# Target engagement studies and kinetic live-cell degradation assays enable the systematic characterization of HDAC6 PROTACs

**DOI:** 10.1101/2025.03.31.646177

**Authors:** Maria Hanl, Felix Feller, Irina Honin, Kathrin Tan, Martina Miranda, Linda Schäker-Hübner, Nico Bückreiß, Matthias Schiedel, Michael Gütschow, Gerd Bendas, Finn K. Hansen

## Abstract

Histone deacetylase 6 (HDAC6) is an important drug target for the treatment of cancer, inflammation, and neurodegenerative disorders. In recent years, the development of proteolysis-targeting chimeras (PROTACs) has emerged to achieve the chemical knockdown of HDAC6. Consequently, there is an urgent need to develop efficient methods for target engagement studies and to enable a thorough characterization of the degradation efficiency and kinetics of HDAC6 PROTACs. In this work, we present a simple NanoBRET assay to assess HDAC6 cellular target engagement using a HeLa^HDAC6-HiBiT^ cell line that stably expresses the LgBiT protein. For this purpose, we successfully designed, synthesized, characterized, and utilized the cell permeable TAMRA-based fluorescent ligand **5**. The key advantage of this NanoBRET assay using HeLa^HDAC6-HiBiT^ cells is the endogenously tagged HDAC6, allowing us to study binding of inhibitors in a near-native environment. Furthermore, we succeeded in establishing a system for kinetic live cell monitoring of HDAC6 degradation. The analysis of the degradation kinetics of a set of HDAC6 PROTACs provided detailed insights into their degradation efficiency and will be helpful for the development of improved HDAC6 degraders in the future.

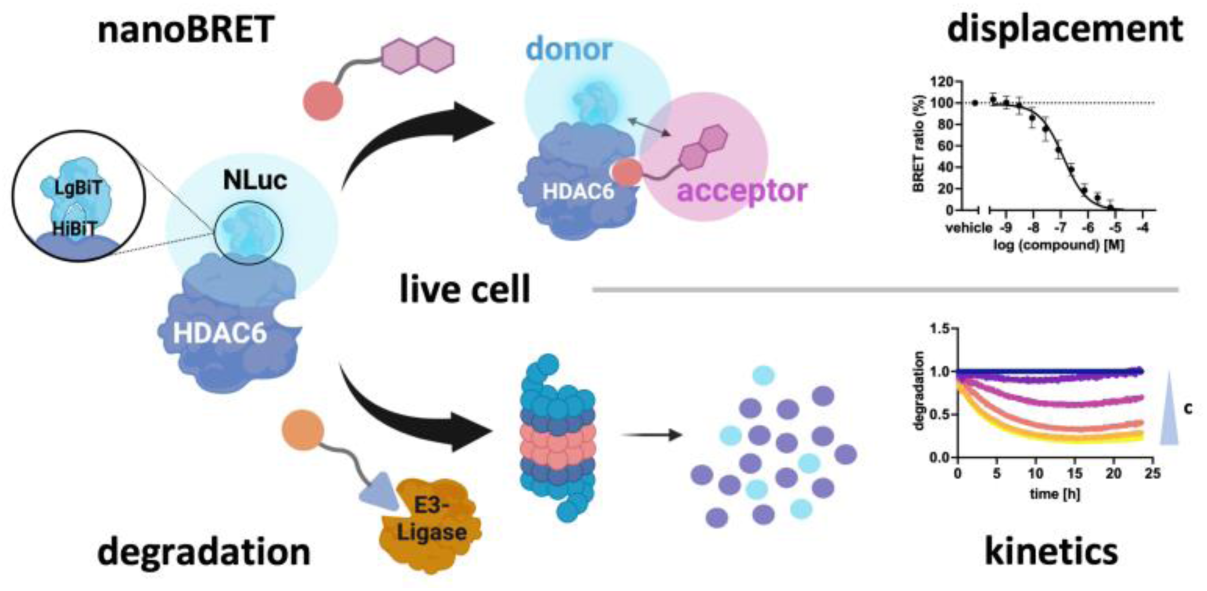

---

Protein acetylation regulates the cellular functions of proteins in the nucleus and cytoplasm and is modulated by histone acetyltransferases (HATs) and histone deacetylases (HDACs).^1^ HDACs catalyze the deacetylation of ε-amino groups of lysine residues in both histone and non-histone proteins. 18 HDACs have been identified in humans and are classified into four classes based on their sequence homology: class I (HDAC1, HDAC2, HDAC3 and HDAC8), class II (IIa: HDAC4, HDAC5, HDAC7, and HDAC9, IIb: HDAC6 and HDAC10), class III (SIRT1, SIRT2, SIRT3, SIRT4, SIRT5, SIRT6, and SIRT7) and class IV (HDAC11). Class I, II, and IV are Zn^2+^-dependent enzymes, while class III isoforms require NAD^+^ as a cofactor.^2–4^

HDAC6 plays a unique role within the HDAC family. Due to its predominant cytoplasmic localization, HDAC6 exhibits distinct substrate specificity, targeting non-histone proteins such as Hsp90, α-tubulin, HSF-1, peroxiredoxin, and cortactin. HDAC6 is involved in numerous physiological cellular processes, such as protein trafficking, angiogenesis, inflammation, and cell migration.^5,6^ Dysregulation of HDAC6 activity is associated with neurodegenerative diseases, various types of cancer, and pathological autoimmune responses.^7–11^

HDAC6 has unique structural characteristics, including two enzymatically active catalytic domains (CD1 and CD2) (Figure 1A). While the function of CD1 remains poorly understood, the CD2 is well characterized for its role in deacetylation.^12^ A dynein motor binding domain (DMB) between these domains facilitates cargo transport along microtubules.^6^ In addition, a ubiquitin-binding zinc finger domain (ZnF-UBP) at the C-terminus promotes the clearance of misfolded proteins.^9,13,14^

**Figure 1.**
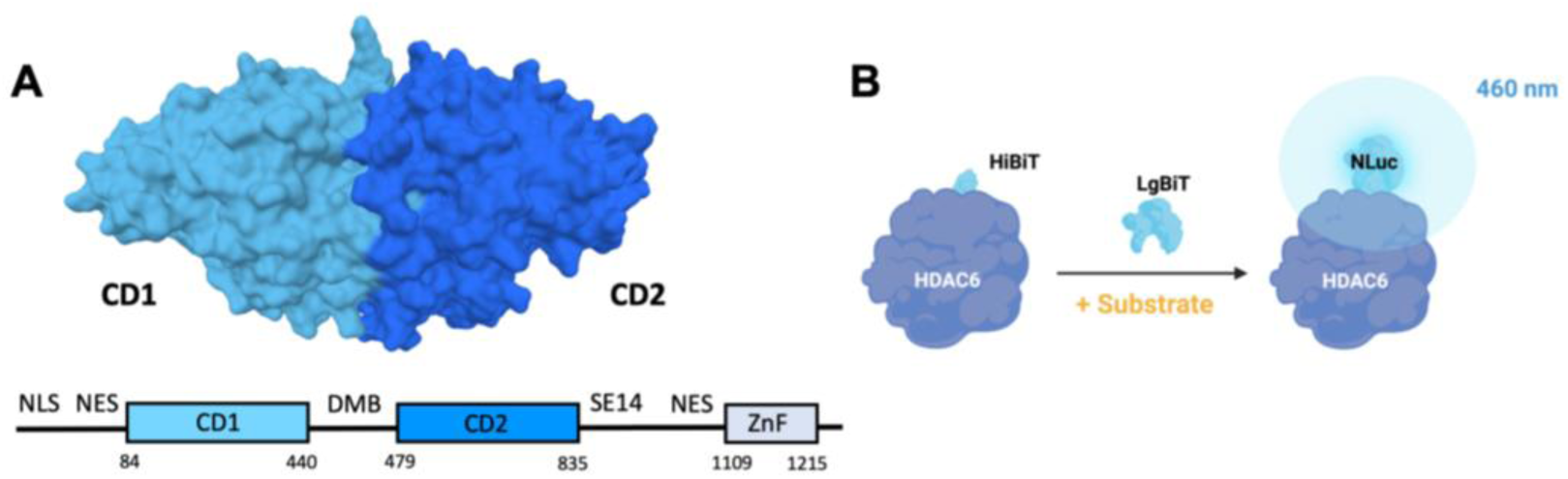
**A**: Illustration of HDAC6 (PDB: 7QNO, from *Danio rerio*). NLS: nuclear localization signal, NES: nuclear export signal, DMB: dynein motor binding, SE14: serine-glutamate tetradecapeptide repeat, ZnF: ubiquitin-binding zinc finger binding domain, CD1: catalytic domain 1, CD2: catalytic domain 2. **B**: Schematic illustration of the HiBiT principle. The active NLuc is formed through binding of the LgBiT to the HiBiT subunit. After the addition of a substrate, a bioluminescence signal can be detected.

To date, five HDAC inhibitors (HDACi) with limited selectivity have been approved by the U.S. Food and Drug Administration (FDA) and the European Medicines Agency (EMA): vorinostat, belinostat, panobinostat, romidepsin, and givinostat. The combination of HDACi with other chemotherapeutic agents or targeted therapies has demonstrated potential as a therapeutic strategy in the treatment of cancer.^15,16^ However, non-selective HDACi are frequently associated with severe adverse effects such as nausea, fatigue, and hematologic toxicities.^17–21^ Thus, selective inhibitors are expected to result in fewer side effects. For instance, HDAC6-preferential inhibitors, such as ricolinostat or citarinostat, are currently in clinical trials, and more recently, highly selective HDAC6 inhibitors have entered clinical trials.^22^

HDAC6 activity can be inhibited either by HDAC6 inhibitors or by protein knockdown using the proteolysis-targeting chimera (PROTAC) technology.^23^ Targeted protein degradation (TPD) *via* PROTACs is an emerging strategy for innovative drug discovery that may offer distinct advantages over classical occupancy-driven inhibitors.^8,24^ Several PROTACs for diverse targets, e.g. vepdegestrant (estrogen receptor), bavdegalutamide (androgen receptor), and CFT8634 (BRD9), have entered clinical trials in recent years.^25–27^

Classical drug design is based on well-defined active sites suitable for small molecules with appropriate pharmacological effects.^28^ However, the design of efficient PROTACs is considerably more complex. PROTACs act through the ubiquitin-proteasome system (UPS), hijacking the intrinsic proteasomal pathway and resulting in a cellular target protein knockdown. PROTACs are typically designed as heterobifunctional molecules consisting of two ligands: one that targets the protein of interest (POI) and another that binds to an E3 ligase, connected by a suitable linker.^29,30^ Protein degradation *via* PROTACs is a multistep, event-driven process coordinated by multiple factors. Once they have reached the target proteins, PROTACs must form binary complexes with the POI or the E3 ligase. These binary complexes facilitate the formation of a stable POI:PROTAC:E3 ligase ternary complex, which promotes polyubiquitination. Subsequently, the polyubiquitinated POI is recognized and degraded by the proteasome.^25,28,31^ A key pharmacological advantage of PROTACs is that they facilitate degradation at sub-stoichiometric concentrations relative to the POI.^31^ Binding affinity does not necessarily correlate with degradation potency; even PROTACs possessing ligands with low affinity can still exert efficient degradation.^23,32,33^ Multiple factors influence the efficacy of PROTACs. For instance, high-affinity binary complexes with a PROTAC can lead to a hook effect at high concentrations, where unproductive binary complexes prevent ternary complex formation.^34^ In contrast, low-affinity binary complexes can sometimes lead to strong degradation.^35^ Moreover, the choice of linker length and flexibility also influences degradation outcomes.^28,34^ It remains a challenge to predict which molecular features are responsible for the formation of a cooperative, stable ternary complex and efficient degradation.^36,37^

The design and optimization of PROTACs is often based on empirical data and extensive chemical optimization.^38^ To better understand these processes in living cells while increasing the predictability of PROTAC activity, it is crucial to develop suitable assay systems. The study of drug-target engagement in living cells is an essential step in drug development.^39^ Therefore, we aimed to develop a NanoBRET assay to investigate the formation of HDAC6:PROTAC binary complexes in an endogenous cellular environment. Traditionally, cellular HDAC6 target engagement studies have relied on antibody-based detection methods such as Western blot, particularly for the detection of tubulin acetylation. Due to its low dynamic range, immunoblot analysis has limited sensitivity in detecting subtle changes in protein acetylation, while tubulin acetylation is influenced not only by HDAC6 but also other factors, including Sirt2, α-tubulin *N*-acetyltransferase (ATAT1), and oxidative stress.^40^ Recently, the first NanoBRET HDAC6 target engagement assays using transfected cells overexpressing the nanoLuc-HDAC6 fusion gene were reported.^41,42^ Nanoluciferase-based systems can be used to track protein levels and study protein interactions in living cells.^41,43–46^ In NanoBRET target engagement assays, binding studies are typically conducted using cells that overexpress the target protein. However, perturbing the natural stoichiometry may cause problems such as aggregation and altered functions.^47,48^ Furthermore, overexpression can affect protein localization and behavior, potentially leading to confounding results when studying protein-protein interactions, protein-ligand interactions, or protein dynamics under natural conditions. To mitigate this issues, it would be advantageous to use the endogenous promotor of the protein and maintain its natural expression level.^37,47^ Schwinn *et al.* developed the HiBiT-LgBiT complementation technology, which enables the investigation of POIs in a native environment (Figure 1B).^47^ The reconstituted nanoluciferase complex (Nluc) consists of an 11-amino acid tag (HiBiT) inserted at the C- or N-terminus of the POI by CRISPR/Cas9 gene editing, and a larger subunit of 18 kDa (termed LgBiT). Co-expression of the POI-HiBiT and LgBiT within the same cell allows for the characterization of target engagement in a near-native environment.

Our group recently developed several PROTACs based on HDAC6 inhibitors.^23,49,50^ In this work, we have therefore focused on establishing methods to investigate intracellular binary complex formation between HDAC6 and PROTACs. Moreover, we have developed a method to precisely characterize the kinetics of PROTAC-induced HDAC6 degradation. Herein, we present a straightforward NanoBRET assay for studying cellular target engagement in a near-native environment, along with the development of an HDAC6 live cell kinetic degradation assay.

## RESULTS AND DISCUSSION

### Design, synthesis, and characterization of a fluorescent probe for HDAC6

For the development of a new NanoBRET assay for HDAC6 we first focused on the design and synthesis of suitable fluorescently labeled tracers. In previous research on NanoBRET tracers for chemokine receptors, we have successfully utilized cell-permeable TAMRA-based fluorophores, which are commonly used as acceptor fluorophores in NanoBRET assays.^51–55^ We planned to introduce the TAMRA fluorophore by a Cu(I) catalyzed Huisgen 1,3-cycloaddition. To verify an appropriately positioned triazole as a suitable connecting unit between the HDACi and TAMRA, we conducted docking studies using the crystal structure of human HDAC6 (PDB: 5EDU). We utilized the hydroxamic acid substructure of vorinostat from the co-crystal structure of *Danio rerio* HDAC6 (PDB: 5EEI) as a template. This substructure was aligned with the human HDAC6 crystal structure to achieve a proper complexation of the zinc ion. Our docking studies with the ligand−triazole conjugate **1** (Figure 2A) indicated a binding mode similar to the parent ligand vorinostat (Figure 2A), i.e. occupying the L1 loop. Additionally, the ethyl-substituted triazole appears to be sufficiently solvent-exposed to serve as an exit vector for fluorophore installation.

**Figure 2.**
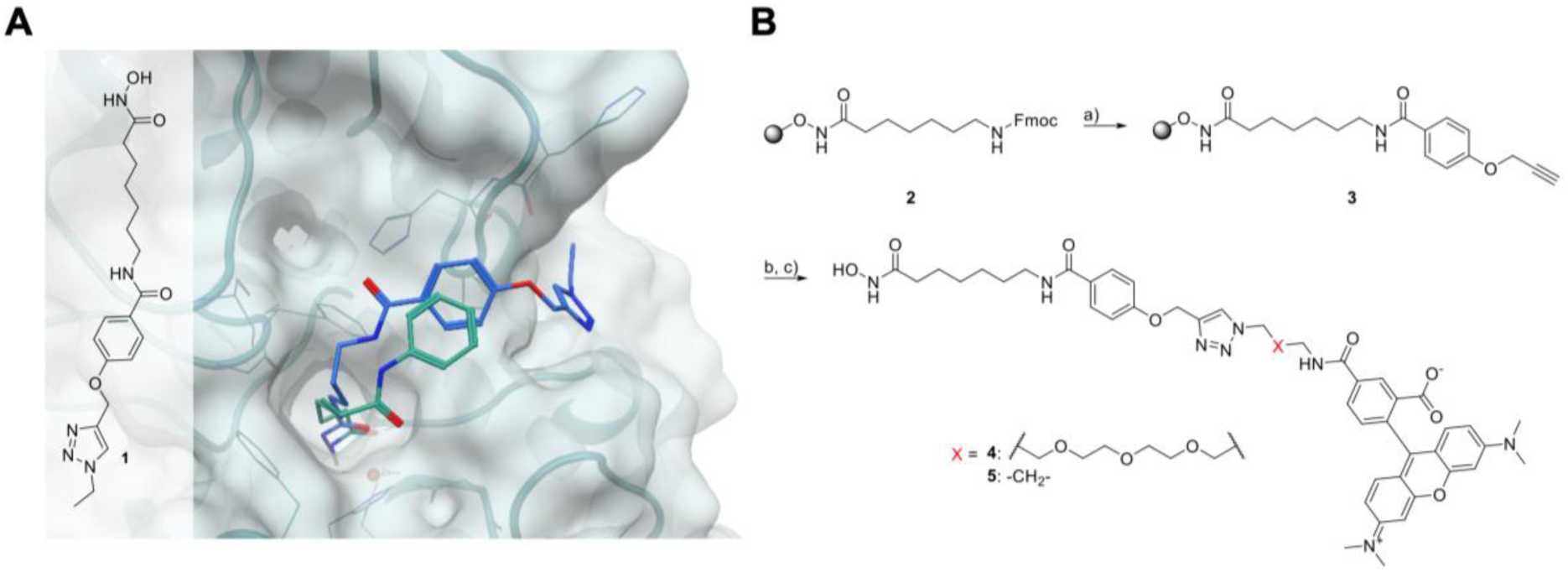
Development of fluorescent ligands for the HDAC6 NanoBRET assay. **A**: Predicted binding mode of the ligand−triazole conjugate **1** (blue colored) in human HDAC6 (PDB: 5EDU). X-ray conformation of vorinostat (turquoise colored) from *Danio rerio* HDAC6 (PDB: 5EEI) placed in human HDAC6 (PDB: 5EDU) *via* superposition. **B**: Synthesis of the TAMRA-based fluorescent HDAC ligands. *Reagents and conditions*: loading of resin **2**: 0.687 mmol/g; (a) (i) 20% piperidine, DMF, rt, 2 x 5 min, (ii) 4-(prop-2-yn-1-yloxy)benzoic acid, HATU, HOBt*H_2_O, DIPEA, DMF, rt, 20 h; (b) azido-functionalized TAMRA analog, TBTA, CuSO_4_*5H_2_O, ascorbic acid, *t*-BuOH, DMF, rt, 18 h; (c) 5% TFA, 5% TIPS, CH_2_Cl_2_, rt, 1 h.

Since the linker length and type can influence cell permeability and fluorescence properties, we designed two different NanoBRET tracers. Compound **4** contains a PEG linker between TAMRA and the triazole connecting unit, while tracer **5** utilizes a shorter propylene linker. The synthesis of both tracers started from our previously published preloaded solid-phase resin HAIR D^56^ (**2**) and is summarized in Figure 2B. In the first step, the Fmoc-protecting group was removed by treatment with piperidine. Subsequently, the HDACi cap was installed using a HATU-mediated amide coupling reaction, yielding the resin-bound intermediate **3** with a *para*-propargyloxy handle. The synthesis of the desired tracers was finalized by performing an on-resin Cu(I)-catalyzed Huisgen cycloaddition reaction with the respective azido-functionalized TAMRA analog, followed by cleavage from the solid support. Both tracers were further purified by preparative RP-HPLC to >95% purity.

### Development of a NanoBRET HDAC6 target engagement assay

To evaluate the inhibitory activity and specificity of the fluorescently labeled ligands **4** and **5** for HDAC6, we initially characterized their binding properties in fluorogenic assays. The biochemical HDAC inhibition assays confirmed that tracers **4** and **5** bind to HDAC1, HDAC2, and HDAC6 with both exhibiting the greatest inhibitory activity against HDAC6 (**4**: IC_50_ = 0.018 µM, **5:** IC_50_ = 0.0091 µM). The inhibitory activity against HDAC6 exceeded that of vorinostat (IC_50_ = 0.034 µM) (Table 1).

**Table 1.** In vitro inhibition of HDAC1, HDAC2, and HDAC6 by tracers 4 and 5.

| | $IC_{50} \pm SD$<br>($\mu M$)<sup>a</sup> | | |
| --- | --- | --- | --- |
|  | HDAC1 | HDAC2 | HDAC6 |
| **4** | $0.20 \pm 0.02$ | $0.34 \pm 0.04$ | $0.018 \pm 0.001$ |
| **5** | $0.13 \pm 0.02$ | $0.22 \pm 0.01$ | $0.0091 \pm 0.0001$ |
| **vorinostat** | $0.15 \pm 0.02$ | $0.25 \pm 0.01$ | $0.034 \pm 0.003$ |

<sup>a</sup>Mean ± SD, duplicate measurement, $n \geq 2$ .

In the next step, we turned our attention towards the development of the NanoBRET-based binding assay. To this end, we first compared the HDAC6 expression levels between HeLa wild-type cells and the commercially available HeLa^HDAC6-HiBiT^ cell line stably expressing the LgBiT protein. Immunoblotting revealed that the HiBiT modification did not cause overexpression of HDAC6. Instead, it slightly reduced the total HDAC6 levels in the modified cell line compared to the wild-type control (Figure S1, Supporting Information).

Afterwards, we developed a NanoBRET-based binding assay (see Figure 3A for a schematic representation). To this end, we investigated the affinity of tracers **4** and **5** (see Figure 3B for extinction and emission spectra of **5**) for HDAC6 in HeLa^HDAC6-HiBiT^ cells stably expressing the LgBiT protein. In whole cell assays, both tracers showed *K*_d_ values in the sub-micromolar concentration range (**4**: *K*_d_ = 0.12 ± 0.04 µM, **5**: *K*_d_ = 0.080 ± 0.011 µM, Figure 3C and D). To assess cell permeability, we permeabilized cells with digitonin and found a lower *K*_d_ value for **4** (*K*_d_ = 0.046 ± 0.021 µM), while the *K*_d_ value for **5** remained unchanged (*K*_d_ = 0.080 ± 0.011 µM). Since tracer **5** demonstrated higher affinity for HDAC6 and favorable cell permeability, we selected it for the subsequent displacement experiments.

**Figure 3.**
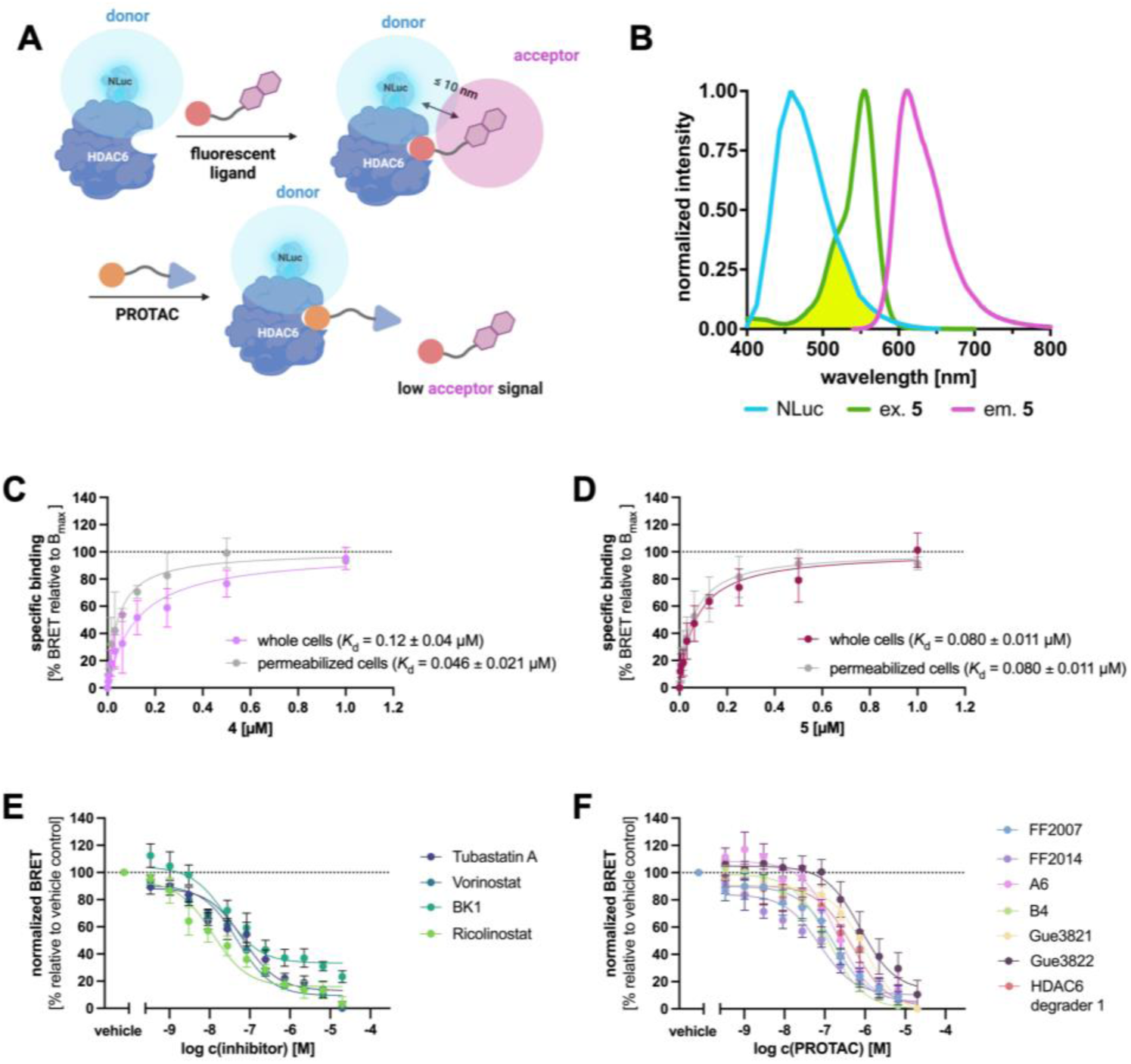
Development of a novel NanoBRET assay for HDAC6 target engagement studies in HeLa^HDAC6-HiBiT^ cells stably expressing the LgBiT protein. **A**: Schematic presentation of the NanoBRET strategy to detect the binding to HDAC6 in cells. **B**: Bioluminescence spectrum of NLuc (blue line) and excitation (green) and emission (pink) spectra of the TAMRA-labeled inhibitor **5**. Energy transfer from NLuc to the tracer fluorophore is detectable through its emission spectrum. The data are normalized to the respective maximum of each spectrum. **C**-**D**: Specific saturation binding curves of **4** (**C**) and **5** (**D**) in a NanoBRET assay in whole cells and after permeabilization with digitonin (50 µg/mL) (mean ± SEM, duplicate measurement, n ≥ 3). **E**-**F**: Competition binding curves of inhibitors (**E**) and PROTACs (**F**) obtained with **5** (500 nM) in whole cells normalized to vehicle control and unspecific binding (mean ± SEM, duplicate measurement, n ≥ 3). Compounds Gue3821 and HDAC6 degrader 1 exhibit autofluorescence at higher concentrations; curves were adjusted to unspecific binding.

In a third step, we evaluated compound **5** as a fluorescent probe to assess HDAC6 target engagement in whole cells. We evaluated a set of HDAC inhibitors (vorinostat, ricolinostat, tubastatin A, and BK1) and recently published PROTACs in competition binding experiments (Figure 4).^23,50,57,58^ Vorinostat is a pan-HDAC inhibitor, ricolinostat preferentially targets HDAC6, tubastatin A is a dual HDAC6/HDAC10 inhibitor, and BK1 is a highly selective HDAC6 inhibitor.^4,59,60^ For the PROTACs, compounds were selected to represent a diverse range of scaffolds, including both non-selective vorinostat-like and selective HDAC6-targeting warheads. Two types of zinc-binding groups (ZBGs) were incorporated: hydroxamic acids and difluoromethyl-1,3,4-oxadiazoles (DFMOs). The E3 ligase ligands used were well-established recruiters of cereblon (CRBN) and the von Hippel–Lindau (VHL) protein.

**Figure 4.**
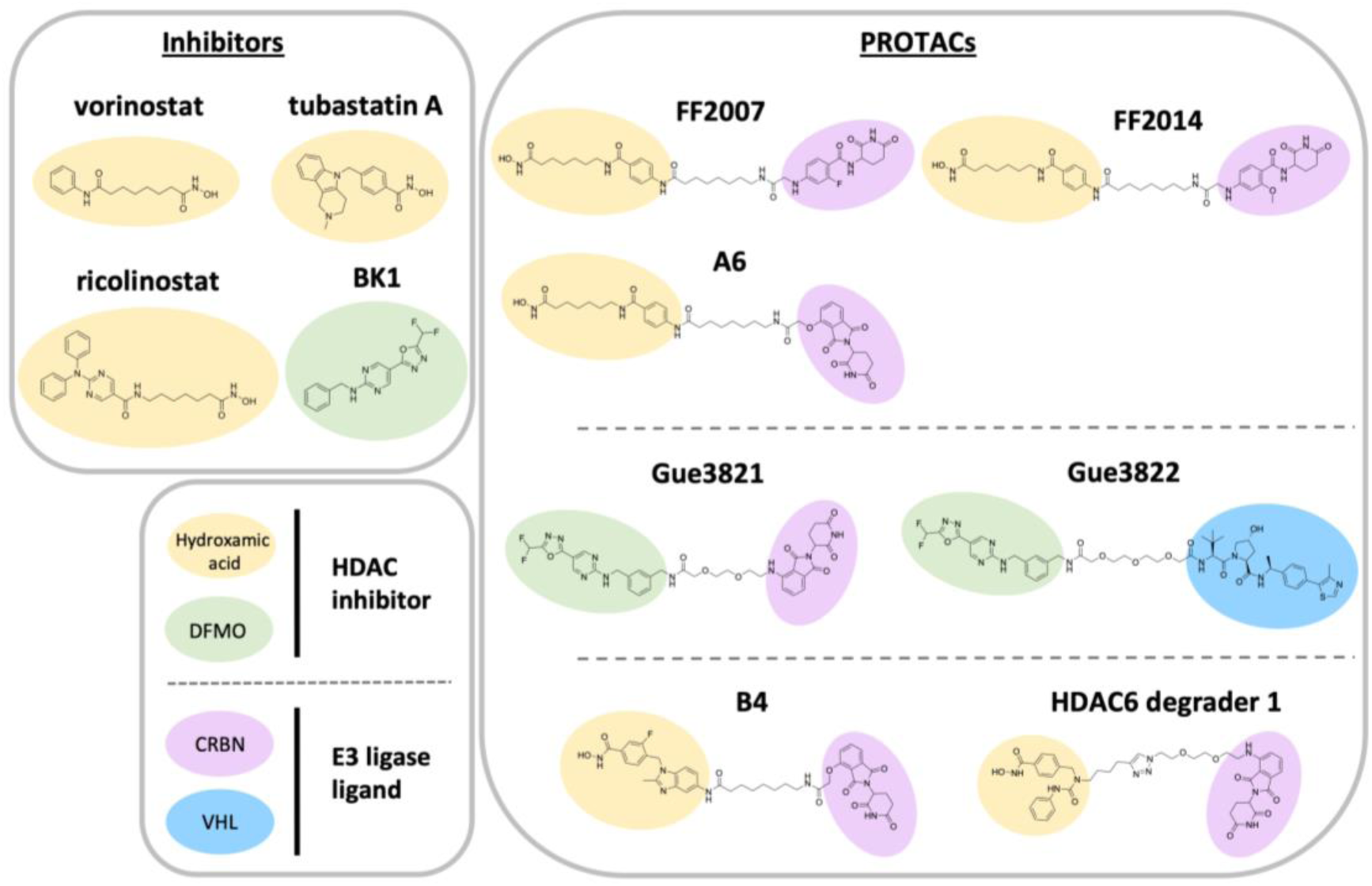
Structures of HDAC inhibitors and PROTACs used in this study. DFMO: difluoromethyl-1,3,4-oxadiazole, CRBN: cereblon, VHL: von Hippel-Lindau.

For the displacement assays, tracer **5** was used at a concentration of 500 nM. The results are shown in Figure 3E, F and Table 2. Compounds Gue3821 and HDAC6 degrader 1 exhibited autofluorescence at higher concentrations (≥ 2.34 µM), which interfered with the NanoBRET signal, as previously reported for other pomalidomide derivatives.^49,61^ Therefore, the calculated IC_50_ values for these compounds represent an upper estimate. Among the HDAC inhibitors, ricolinostat exhibited the highest potency in displacing tracer **5** (IC_50_ = 0.021 ± 0.011 µM). Surprisingly, tubastatin A showed the lowest potency in the NanoBRET assay (IC_50_ = 0.091 ± 0.024 µM), but demonstrated stronger target engagement than previously reported in an overexpressing cell system using a different fluorescent tracer.^41^ Notably, tubastatin A exhibited the highest potency in the biochemical HDAC6 inhibition assay (Table 2).

**Table 2.** NanoBRET binding results obtained with tracer 5 (500 nM).

|  | HDAC6<br>enzyme inhibition | NanoBRET assay<sup>a</sup> |  |
| --- | --- | --- | --- |
|  | IC<sub>50</sub> ± SD<br>(μM) | IC<sub>50</sub> ± SEM<br>(μM) | K<sub>i</sub> ± SEM<br>(μM) |
| **tubastatin A** | 0.014 ± 0.001<sup>62</sup> | 0.091 ± 0.024 | 0.011 ± 0.004 |
| **vorinostat** | 0.034 ± 0.003 | 0.046 ± 0.008 | 0.0057 ± 0.0010 |
| **BK1** | 0.19 ± 0.01<sup>60</sup> | 0.075 ± 0.030 | n.d.<sup>c</sup> |
| **ricolinostat** | 0.018 ± 0.003<sup>62</sup> | 0.021 ± 0.011 | 0.0015 ± 0.0008 |
| **FF2007** | 0.0068 ± 0.0005 | 0.17 ± 0.04 | 0.025 ± 0.005 |
| **FF2014** | 0.0051 ± 0.0009 | 0.14 ± 0.06 | 0.015 ± 0.004 |
| **A6** | 0.0047 ± 0.0003<sup>23</sup> | 0.23 ± 0.06 | 0.030 ± 0.008 |
| **B4** | 0.0045 ± 0.0004<sup>23</sup> | 0.15 ± 0.05 | 0.021 ± 0.008 |
| **Gue3821** | 0.64 ± 0.21<sup>50</sup> | 1.0 ± 0.2<sup>b</sup> | n.d.<sup>c</sup> |
| **Gue3822** | 0.69 ± 0.12<sup>50</sup> | 0.82 ± 0.16 | n.d.<sup>c</sup> |
| **HDAC6 degrader 1** | 0.026 ± 0.002 | 0.59 ± 0.21<sup>b</sup> | 0.083 ± 0.030<sup>b</sup> |

<sup>a</sup>IC<sub>50</sub> values were calculated from competition binding curves (mean ± SEM, duplicate measurement, n ≥ 3). <sup>b</sup>Compounds showing autofluorescence at higher concentrations; curves were adjusted to unspecific binding. <sup>c</sup>n.d.: not determined, K<sub>i</sub> values were calculated for the compounds that fulfill the criteria defined by the Cheng-Prusoff equation.

The results for the PROTACs are presented in Figure 3F and Table 2. Structurally, some PROTACs share common features, including similar scaffolds and linkers. Compounds FF2007, FF2014, and A6 have the same vorinostat-like HDAC scaffold and a cognate linker.^23,57^ Interestingly, A6 displayed a higher IC_50_ value (IC_50_ = 0.23 ± 0.06 µM) in the NanoBRET assay than FF2007 and FF2014, although these compounds evoke similar HDAC6 inhibition in the enzymatic assay.

Compounds Gue3821 and Gue3822, both based on the slow-binding selective HDAC6 inhibitor BK1^50^ utilizing a difluoromethyl-1,3,4-oxadiazole (DFMO) ZBG, exhibited higher IC_50_ values in the displacement assays (Gue3821: IC_50_ = 1.0 ± 0.2 µM, Gue3822: IC_50_ = 0.82 ± 0.16 µM). The well-established reference PROTAC HDAC6 degrader 1^8^ showed a moderate IC_50_ value in the NanoBRET assay (IC_50_ = 0.59 ± 0.21 µM) and also in the enzymatic HDAC6 inhibition assay. To determine whether the reduced IC_50_ values of the PROTACs observed in the NanoBRET assay were due to limited cell permeability, we conducted a cell permeability study using a cellular HDAC inhibition assay in both intact and lysed HeLa^HDAC6-HiBiT^ cells. Comparison of the PROTACs FF2014 and A6 with the inhibitor vorinostat showed that FF2014 and A6 had reduced cell permeability, whereas vorinostat did not (Figure S2, Supporting Information). These findings suggest that the discrepancies observed for the PROTACs between biochemical assays using isolated enzymes and the cellular NanoBRET assay likely result from limited cellular permeability, a common limitation of PROTACs due to their high molecular weight.^63^

In addition to the IC_50_ values for the HDAC inhibitors and the PROTACs, we also calculated the *K*_i_ values for the compounds meeting the criteria defined by the Cheng-Prusoff equation (Table 2).^64^ Unlike the IC_50_ values from the displacement assay, which are highly dependent on assay conditions, the equilibrium dissociation constant (*K*_i_) values calculated using the Cheng-Prusoff equation are system independent. The tracer concentration present in the medium was used for the calculation of *K*_i_. Since BK1 inhibits HDAC6 through a distinct two-step slow binding mechanism that falls outside the criteria defined by the Cheng-Prusoff equation, this may also apply for the derived PROTACs (Gue3821 and Gue3822). Therefore, *K*_i_ values were not calculated for these compounds.

Our results from the NanoBRET target engagement assay with endogenously tagged HDAC6 confirmed the NanoBRET assay to be a suitable method for studying target engagement and binary complex formation between HDAC6 and PROTACs. Furthermore, these results suggest that the enzymatic assay may not necessarily be a reliable indicator of target engagement potency in a native environment.

### Development of a kinetic live-cell HDAC6 degradation assay

To establish the HiBiT live-cell kinetic system for characterizing our set of HDAC6-targeting PROTACs (Figure 4) in a native environment, and to detect HDAC6 degradation (see Figure 5A for a schematic representation of the assay principle), we first conducted an initial screening using immunoblotting analysis. This screening was performed in HeLa^HDAC6-HiBiT^ cells stably expressing the LgBiT protein and MM.1S cells (Figure 5B). In the next step, we evaluated the HDAC6 degradation using the lytic HiBiT detection assay in HeLa^HDAC6-HiBiT^ cells stably expressing the LgBiT protein (Figure 5C). Western blot analysis revealed higher HDAC6 degradation in MM.1S cells compared to HeLa^HDAC6-HiBiT^ cells, suggesting the degradation efficiency to be cell line-dependent. Although the less efficient degradation was observed in HeLa^HDAC6-HiBiT^ cells, a strong correlation was found between the results of the immunoblot analysis and the lytic HiBiT detection assay (24 h: r^2^ = 0.98, slope = 1.03) in HeLa^HDAC6-HiBiT^ cells (Figure S3, Supporting Information).

**Figure 5.**
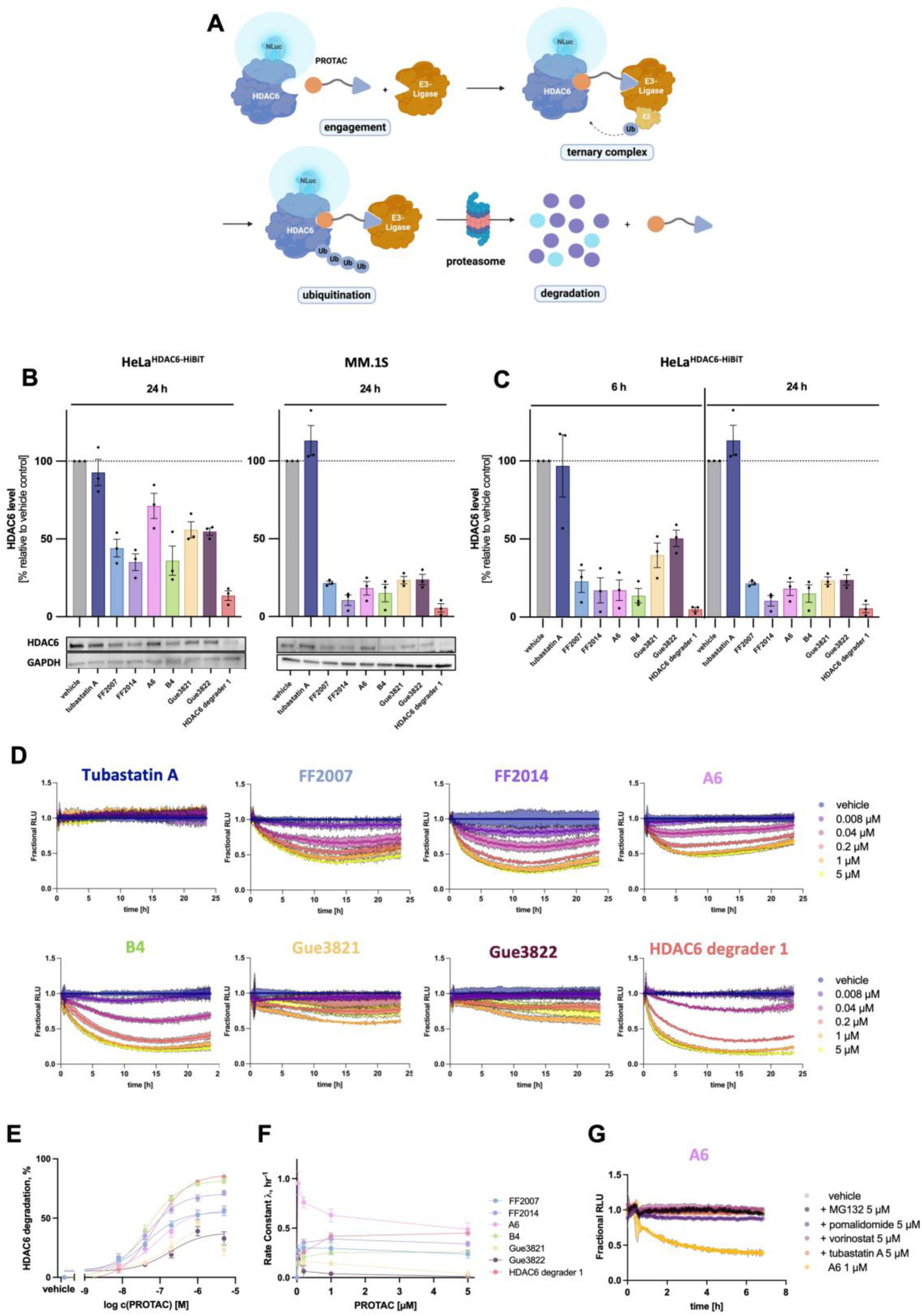
Development of a novel HiBiT live-cell kinetic degradation assay. **A**: Schematic presentation of the HiBiT strategy to detect the degradation of HDAC6 in cells. **B**-**C**: Screening of HDAC6 degradation after treatment with the PROTACs at 1 µM. **B**: Western Blot with HeLa^HDAC6-HiBiT^ cells stably expressing the LgBiT protein and MM.1S cells with 1 µM of PROTACs after 24 h incubation with a representative image. GAPDH was used as loading control (mean ± SEM, n = 3). **C**: HiBiT lytic detection with HDAC6 target engagement in HeLa^HDAC6-HiBiT^ cells stably expressing the LgBiT protein with 1 µM after 6 h and 24 h incubation (mean ± SEM, triplicate measurement, n = 3). **D**: Live cell HDAC6 degradation kinetics for various PROTACs were assessed using the HiBiT live-cell degradation kinetic assay in HeLa^HDAC6-HiBiT^ cells stably expressing the LgBiT protein. PROTACs were applied at the indicated concentrations for 24 hours (mean ± SEM, triplicate measurement, n ≥ 3). RLU: relative light units. **E**-**F**: Determination of the Dmax_50_ value (**E**), and degradation rate (**F**) (mean ± SEM, triplicate measurement, n ≥ 3). **G**: Rescue experiments were performed with MG132, pomalidomide, vorinostat, and tubastatin A at the indicated concentrations, using A6 as the representative PROTAC (mean ± SD, triplicate measurement, n = 2).

Next, we monitored the degradation kinetics for HDAC6 using the HiBiT live-cell kinetic assay in HeLa^HDAC6-HiBiT^ cells stably expressing LgBiT. The kinetic data are presented in Figure 5D-F and are consistent with the results from the immunoblot and the lytic HiBiT detection assays. Cell viability was assessed at the endpoint of the kinetic assay to rule out potential toxicity issues (Figure S4, Supporting Information).

To better understand the differences in degradation kinetics among the PROTACs (see Table 3 for degradation rates, Dmax and Dmax_50_ values, see Experimental Section for details), we examined their structural features in detail. First, we inspected compounds Gue3821 and Gue3822, both based on the slow-binding DFMO-based inhibitor BK1. Although Gue3821 (a CRBN recruiter) showed slightly more efficient degradation than Gue3822 (a VHL recruiter), both compounds exhibited the weakest HDAC6 degradation among this set of PROTACs. Their degradation followed almost linear kinetics, with a hook effect observed at 5 µM. At higher concentrations, the degradation rates of both Gue3821 and Gue3822 also decreased, likely due to the hook effect. DFMO-based inhibitors such as BK1 exhibited a two-step slow binding mechanism, which may also apply to Gue3821 and Gue3822.^60,65^

**Table 3.** Calculation of the descriptive parameters of the HDAC6 degradation kinetics.

| | Parameter | Concentration ($\mu\text{M}$)<sup>a</sup> | | | | | Dmax<sub>50</sub><br>($\mu\text{M}$)<sup>a</sup> |
| --- | --- | --- | --- | --- | --- | --- | --- |
|  |  | 0.008 | 0.04 | 0.2 | 1 | 5 |  |
| FF2007 | Rate ($\text{hr}^{-1}$) | n.d.<sup>b</sup> | $0.27 \pm 0.07$ | $0.30 \pm 0.05$ | $0.30 \pm 0.06$ | $0.24 \pm 0.04$ | $0.042 \pm 0.006$ |
| | Dmax (%) | $16 \pm 3$ | $28 \pm 5$ | $44 \pm 5$ | $53 \pm 5$ | $56 \pm 5$ | |
| | AUC | $22 \pm 1$ | $19 \pm 2$ | $16 \pm 1$ | $14 \pm 1$ | $14 \pm 1$ | |
| FF2014 | Rate ($\text{hr}^{-1}$) | n.d.<sup>b</sup> | $0.28 \pm 0.02$ | $0.35 \pm 0.04$ | $0.39 \pm 0.04$ | $0.34 \pm 0.03$ | $0.043 \pm 0.006$ |
| | Dmax (%) | $17 \pm 4$ | $35 \pm 4$ | $57 \pm 3$ | $67 \pm 3$ | $71 \pm 3$ | |
| | AUC | $21 \pm 1$ | $18 \pm 1$ | $13 \pm 1$ | $11 \pm 1$ | $9.8 \pm 0.6$ | |
| A6 | Rate ($\text{hr}^{-1}$) | n.d.<sup>b</sup> | $1.0 \pm 0.2$ | $0.76 \pm 0.06$ | $0.63 \pm 0.07$ | $0.49 \pm 0.07$ | $0.085 \pm 0.028$ |
| | Dmax (%) | $11 \pm 3$ | $24 \pm 4$ | $38 \pm 5$ | $52 \pm 4$ | $57 \pm 4$ | |
| | AUC | $23 \pm 1$ | $20 \pm 2$ | $17 \pm 2$ | $15 \pm 2$ | $13 \pm 1$ | |
| B4 | Rate ($\text{hr}^{-1}$) | n.d.<sup>b</sup> | $0.14 \pm 0.03$ | $0.21 \pm 0.03$ | $0.25 \pm 0.03$ | $0.26 \pm 0.03$ | $0.044 \pm 0.007$ |
| | Dmax (%) | $13 \pm 4$ | $40 \pm 5$ | $66 \pm 3$ | $79 \pm 3$ | $81 \pm 3$ | |
| | AUC | $22 \pm 1$ | $16 \pm 1$ | $11 \pm 1$ | $7.6 \pm 0.3$ | $7.0 \pm 0.2$ | |
| Gue3821 | Rate ($\text{hr}^{-1}$) | n.d.<sup>b</sup> | $0.15 \pm 0.08$ | $0.16 \pm 0.04$ | $0.14 \pm 0.03$ | $0.037 \pm 0.032$ | $0.23 \pm 0.06^c$ |
| | Dmax (%) | $14 \pm 5$ | $12 \pm 2$ | $25 \pm 5$ | $47 \pm 3$ | $23 \pm 5$ | |
| | AUC | $22 \pm 1$ | $22 \pm 1$ | $19 \pm 2$ | $15 \pm 1$ | $20 \pm 1$ | |
| Gue3822 | Rate ($\text{hr}^{-1}$) | n.d.<sup>b</sup> | $0.19 \pm 0.05$ | $0.063 \pm 0.060$ | $0.039 \pm 0.018$ | $0.011 \pm 0.007$ | $0.16 \pm 0.03$ |
| | Dmax (%) | $11 \pm 2$ | $11 \pm 1$ | $19 \pm 3$ | $39 \pm 4$ | $33 \pm 6$ | |
| | AUC | $23 \pm 1$ | $23 \pm 1$ | $21 \pm 1$ | $17 \pm 1$ | $19 \pm 2$ | |
| HDAC6<br>degrader 1 | Rate ($\text{hr}^{-1}$) | n.d.<sup>b</sup> | $0.21 \pm 0.02$ | $0.26 \pm 0.03$ | $0.42 \pm 0.04$ | $0.45 \pm 0.03$ | $0.077 \pm 0.008$ |
| | Dmax (%) | $6.7 \pm 2.9$ | $27 \pm 2$ | $65 \pm 4$ | $80 \pm 2$ | $85 \pm 2$ | |
| | AUC | $23 \pm 1$ | $19 \pm 1$ | $11 \pm 1$ | $7.0 \pm 0.6$ | $5.8 \pm 0.4$ | |

<sup>a</sup>Mean ± SEM, triplicate measurement, $n \geq 3$ . <sup>b</sup>n.d. not determined, fitting not possible. <sup>c</sup>Hook effect
at 5 $\mu\text{M}$ , curve was adjusted to 1 $\mu\text{M}$ .

Next, the two most potent PROTACs B4^23^ and HDAC6 degrader 1 were examined in detail. Interestingly, these compounds are structurally diverse, featuring different HDAC6 inhibitors, linker types, and linker lengths, but both share a similar CRBN recruiter. HDAC6 degrader 1 exhibited the highest potency and the fastest degradation. However, a comparison of the Dmax_50_ revealed a lower Dmax_50_ value for B4 (Dmax_50_ = 0.044 ± 0.007 µM) than for HDAC6 degrader 1 (Dmax_50_ = 0.077 ± 0.008 µM). We then analyzed the group consisting of FF2007, FF2014, and A6. All three compounds are based on the same pan-HDAC inhibitor scaffold and share an identical linker. A6 utilizes pomalidomide as a CRBN recruiter, while FF2007 and FF2014 use benzamide-based CRBN recruiters. Within this group, FF2014 was the most efficient degrader (at 5 µM: Dmax = 71 ± 3%) compared to FF2007 (at 5 µM: Dmax = 56 ± 5%) and A6 (at 5 µM: Dmax = 57 ± 4%). These results were in agreement with previously published data.^57^ The Dmax_50_ values of FF2007 (Dmax_50_ = 0.042 ± 0.006 µM) and FF2014 (Dmax_50_ = 0.043 ± 0.006 µM) were highly similar. Interestingly, A6 showed a slight HDAC6 recovery over time, consistent with findings from a previous report using HiBiT-tagged K562 cells.^23^ At high concentrations, A6 demonstrated a clear trend of a decelerating degradation rate (Figure 5F); however, the HDAC6 degradation provoked by A6 remained the fastest of all PROTACs tested. This decrease was not associated with a hook effect.

Next, we successfully confirmed the dependence of HDAC6 degradation on ternary complex formation and the UPS using A6 as a representative example by co-treatment with MG132, pomalidomide, vorinostat, and tubastatin A (Figure 5G). While A6 alone induced a substantial reduction of HDAC6 protein levels, all co-treatment conditions resulted in a rescue of the A6-induced degradation. These data corroborated that A6-mediated HDAC6 degradation is UPS-dependent and requires binding to CRBN and HDAC6.

Finally, we evaluated whether the kinetic live-cell HDAC6 degradation assay can be used to characterize HDAC degraders featuring alternative E3 ligase ligands and HDAC-binding moieties (Figure 6A). To this end, we selected FF2049,^66^ a Fem-1 homologue B (FEM1B)-recruiting, selective HDAC1–3 degrader that shares the HDAC inhibitor scaffold of FF2007, FF2014, and A6, but employs a covalent chloroacetyl-based warhead to engage FEM1B. Additionally, we designed and synthesized MM-47 (see Experimental Section for synthetic details), a potential HDAC6 degrader incorporating a phenyl glutarimide-based CRBN ligand and a tubastatin A-derived HDAC6-binding warhead. Since FF2049 showed cytotoxicity at concentration higher than 0.2 µM (see Figure S4, Supporting Information) we only evaluated the kinetic traces up to this concentration. Consistent with previously reported immunoblot results,^66^ FF2049 showed no HDAC6 degradation in the live-cell kinetic assay (Figure 6B). In contrast, MM-47 induced substantial HDAC6 degradation (Dmax_50_ = 0.014 **±** 0.003) (Figure 6C-D). Interestingly, MM-47 demonstrated the slowest degradation rate of all tested hydroxamic acid-based degraders (Figure 6E).

**Figure 6:**
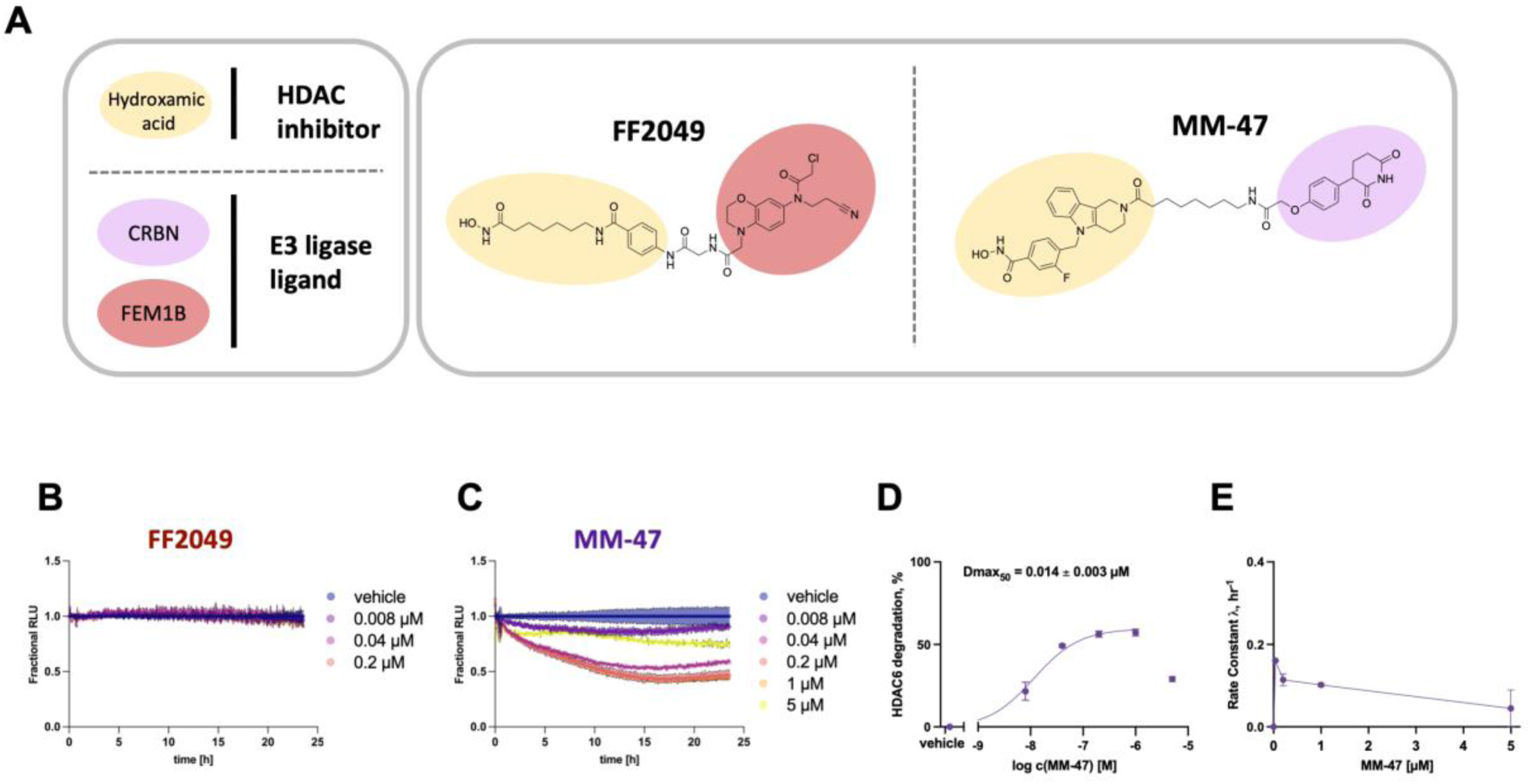
Characterization of PROTACs with alternative E3 ligase ligands and HDAC-binding moieties. **A:** Structures of the PROTACs FF2049 and MM-47. **B-D:** Live cell HDAC6 degradation kinetics using the HiBiT live-cell degradation kinetic assay in HeLa^HDAC6-HiBiT^ cells stably expressing the LgBiT protein for FF2049 (**B**) and MM-47 (**C**). PROTACs were applied at the indicated concentrations for 24 hours (mean ± SEM, triplicate measurement, n = 2). **D-E:** Determination of the Dmax_50_ value (**D**), and degradation rate (**E**) of MM-47 (mean ± SEM, triplicate measurement, n = 2).

Taken together, our results demonstrate that the HDAC6 HiBiT live-cell kinetic system in an endogenous environment is an effective tool for a rapid and in-depth elucidation of the degradation efficiency of PROTACs. The data presented provide valuable guidance for the optimization of structural features to improve PROTAC design targeting HDAC6. Furthermore, our results highlight the limitations of Western blotting, which lacks the sensitivity and dynamic range required to accurately measure subtle differences in the degradation performance of PROTACs.

## CONCLUSION

The lack of robust tools to study cellular HDAC6 target engagement and degradation in a near-native environment has limited the progress in the field of targeted HDAC6 degraders. In this study, we developed tools to investigate HDAC6 target engagement and degradation in living cells. To this end, we applied the NanoBRET assay technology to detect binary complex formation of HDAC6 inhibitors and degraders with HDAC6. This method provides a better understanding of target-PROTAC interaction in near-native environments compared to the previously used approaches. In addition, we have established a live-cell assay to track the degradation of HDAC6 using the HiBiT-LgBiT complementation technology. Unlike traditional Western blot techniques, this kinetic assay overcomes the challenges of time and concentration variability by providing real-time insights. The resulting degradation profiles offer a range of key parameters for multidimensional analysis and provide insights into mechanistic aspects that can guide the optimization of PROTACs. Collectively, our findings will support future investigations on HDAC6 degradation, offer new perspectives on the mechanism of action of PROTACs, and will serve as a promising basis for future medicinal chemistry campaigns to design and discover effective HDAC6 degraders with high specificity and efficiency.

## METHODS

### Biological experiments

#### Cell lines and cell culture

The HDAC6-HiBiT KI HeLa (LgBiT) cell line was purchased from Promega (Promega, Madison, WI, USA, #CS3023361) and used as received. This heterozygous clonal cell line was cultured in DMEM (Life Technologies, Darmstadt, Germany) supplemented with 10% fetal bovine serum (Sigma Aldrich, St. Louis, MO, USA), 100 IU/mL penicillin, and 0.1 mg/mL streptomycin (PAN Biotech GmbH, Aidenbach, Germany). MM.1S cells (ATCC, Manassas, VA, USA, CRL-2974) were cultured in RPMI 1640 medium (Life Technologies,) supplemented with 10% fetal bovine serum (PAN Biotech GmbH,), 100 IU/mL penicillin and 0.1 mg/mL streptomycin (PAN Biotech GmbH,) and 1 mM sodium pyruvate (ThermoFisher Scientific Inc.; Waltham, MA, USA). The cell lines were incubated at 37 °C under humidified air with 5% CO_2_. Cell cultures were regularly monitored for mycoplasma contamination.

#### Live-cell kinetic HiBiT detection

In the following, an automated pipetting robot system (Integra Biosciences, Bibertal, Germany) was used. HeLa^HDAC6-HiBiT^ cells stably expressing the LgBiT protein were plated in white 384-well plates (Greiner Bio-One, Kremsmuenster, Austria) at a density of 8 × 10^3^ cells/well and left to attach at 37 °C and 5% CO_2_ under humidified air for 2 hours or overnight. An initial dilution series of test compounds was prepared in DMSO. Subsequently, the dilution was diluted in CO_2_-independent medium (Life Technologies). The CO_2_-independent medium was supplemented with 10% fetal bovine serum (Sigma Aldrich, St. Louis, MO, USA), 4 mM L-glutamine (PAN Biotech GmbH), 100 IU/mL penicillin, and 0.1 mg/mL streptomycin (PAN Biotech GmbH). The final DMSO concentration was 0.25%. An equal volume of CO_2_-independent medium containing a 2-fold concentration of Nano-Glo® Endurazine^TM^ Live Cell Substrate (Promega, Madison, WI, USA) was added to the cells according to the manufacturer’s protocol and incubated for 2.5 h at 37 °C and 5% CO_2_ under humidified air. Luminescence was then measured. Subsequently, the compounds were added, the plate was sealed with Breath-Easy (Diversified-Biotech, Dedham, MA, USA) and luminescence was measured every 5 minutes for 24 h at 37 °C in a Tecan Spark (Tecan Group AG, Maennedorf, Swiss). The data analysis was performed using Excel. The data were first normalized to the luminescence of each well before PROTAC addition and then normalized to the mean of the replicates from the vehicle control at each time point. Further calculations were performed using Graph Pad Prism (Graph Pad Prism 9.0, San Diego, CA, USA). The degradation rate constant was calculated using only the initial degradation until the data reaches the plateau, utilizing the one phase decay equation. Dmax values were determined as the lowest luminescence reading in each well and represented as a percentage of degradation. Dmax_50_ values were calculated by plotting Dmax against concentration using the 3-parameter logistic equation.

#### Lytic HiBiT detection

An automated robotic pipetting system (Integra Biosciences, Bibertal, Germany) was used for the following steps. HeLa^HDAC6-HiBiT^ cells stably expressing the LgBiT protein were plated in white 384-well plates (Greiner Bio-One) at a density of 8 × 10^3^ cells/well and left to attach at 37 °C and 5% CO_2_ under humidified air for 1-2 h or overnight. An initial dilution series of test compounds was prepared in DMSO. This dilution was then further diluted in medium. The final DMSO concentration was 0.25%. Compounds were added to the cells at a final concentration of 1 µM and incubated for 6 h or 24 h at 37 °C and 5% CO_2_ under humidified air. The 2-fold lytic detection reagent was prepared immediately before use according to the manufacturer’s protocol (Promega, Madison, WI, USA) and added to the cells. The plate was mixed for 10 minutes at 350 rpm. Luminescence was measured with a Tecan Spark (Tecan Group AG). Calculations were performed using Graph Pad Prism (Graph Pad Prism 9.0).

#### Cell viability assay

Compound toxicity was determined directly after the 24 h live cell kinetic HiBiT detection. 20 µL of CellTiter-Glo® 2.0 Cell Viability Assay (Promega, Madison, WI, USA) was added to each well. The plate was shaken at 500 rpm for 5 minutes on an orbital shaker and left for 30 minutes at room temperature. Then, the luminescence was measured. The data were normalized to vehicle control.

#### NanoBRET assay

An automated pipetting robot system (Integra Biosciences) was used. HeLa^HDAC6-HiBiT^ cells stably expressing the LgBiT protein were plated in white 384-well plates with nonbinding surface (PerkinElmer, Waltham, MA, USA) at a density of 2 × 10^4^ cells/well in Opti-MEM® I (Life Technologies). For the dilution series, an initial series dilution was first prepared in DMSO. This dilution was then further diluted in Opti-MEM® I. The final DMSO concentration was 0.25%.

For saturation binding experiments in whole cells, a serial dilution of the tracer was added to the cells. An unlabeled ligand (vorinostat 20 µM) was added to determine non-specific binding. Cells were incubated at 37 °C and 5% CO_2_ for 2 h prior to BRET measurements. For saturation binding experiments with permeabilized cells, digitonin (final concentration 50 µg/mL) was additionally added. These plates were incubated at room temperature in the dark for 30 minutes prior to BRET measurements. For BRET measurements, the NanoBRET™ Nano-Glo® Substrate (Promega, Madison, WI, USA) was diluted in Opti-MEM® I (final assay dilution: 1:500).

For competition binding experiments, a final tracer concentration of 500 nM was used. The dilution series of the substances were added to the cells. Cells were incubated at 37 °C and 5% CO_2_ for 2 h prior to BRET measurements. For BRET measurements, the NanoBRET™ Nano-Glo® Substrate was diluted in Opti-MEM® I (final assay dilution 1:1000).

The NanoBRET™ Nano-Glo® Substrate was added and incubated at room temperature for 2-3 minutes. Luminescence was measured with a Tecan Spark (Tecan Group AG) using 465/20 nm (donor) and 615/20 nm (acceptor) filters. BRET-Ratios were calculated in milliBRET Units (mBU). Data were analyzed using Graph Pad Prism (Graph Pad Prism 9.0). Specific binding was determined by subtracting the non-specific binding from the total binding. Tracer affinity was determined using the equations of one-site saturation binding. Competition binding was analyzed using the 3-parameter logistic equation and the one site-fit *K*_i_ equation.

#### Immunoblot

MM.1S cells (1 × 10^6^ cells/mL) and HeLa^HDAC6-HiBiT^ cells stably expressing the LgBiT protein (0.5 × 10^6^ cells/mL) were seeded to cell culture flasks and treated with 1 µM of compound or vehicle (DMSO) for 6 or 24 h. The final DMSO concentration was 0.01%. Cell lysis was performed with Cell Extraction Buffer (Thermo Fisher Scientific Inc) supplemented with Halt Protease Inhibitor Cocktail (100X) (Life-Technologies GmbH) and phenylmethylsulfonyl fluoride (Sigma-Aldrich) according to the manufacturer’s protocol. The protein content was determined with Pierce^TM^ BCA Protein Assay Kit (Thermo Fisher Scientific Inc.) according to the manufacturer’s protocol. Samples were denatured with Laemmli 2X concentrate (Sigma-Aldrich). Precision Plus Protein Unstained Standard (Bio-Rad, Hercules, CA, USA) was used as standard marker. SDS-PAGE was performed with 4-20% Mini-PROTEAN® TGX Stain-Free Protein Gel (Bio-Rad, Hercules) at 200 V for 30 minutes. Proteins were transferred with Trans-Blot Turbo Mini 0.2 µM PVDF Transfer Packs (Bio-Rad) with the Trans-Blot Turbo Transfer System (Bio-Rad) at 2.5 A for 3 minutes. Then membranes were incubated with blocking buffer for 1 h at room temperature. Afterwards membranes were incubated with 1:1000 anti-HDAC6 (Cell Signaling Technology, Denver, MA, USA) and 1:20,000 anti-GAPDH (Affinity Biosciences, Cincinnati, OH, USA) at 4 °C overnight. After washing, membranes were incubated with HRP-conjugated secondary anti-mouse (Santa Cruz, Dallas, TX, USA) and anti-rabbit (R&D Systems Inc., Minneapolis, MN, USA) antibody solutions for 1.5 h. Membranes were developed with Clarity western ECL substrate (Bio-Rad). Detection was performed with ChemiDox XRS+ System (Bio-Rad). Data quantification was performed with Image Lab Software 6.1 (Bio-Rad). Further analysis was performed with GraphPad Prism (Graph Pad Prism 9.0).

#### HDAC inhibition assay

The *in vitro* HDAC inhibition assay for human HDAC1, HDAC2 and HDAC6 inhibitory activity was conducted following a previously published protocol.^62,67^ The assay was performed in black 96-wells OptiPlates (PerkinElmer, Waltham, MA, USA) with a final assay volume of 100 µL per well. 5.0 µL of the test compound or control, diluted in assay buffer (50 mM Tris-HCl, pH 8.0, 137 mM NaCl, 2.7 mM KCl, 1.0 mM MgCl_2_, 0.1 mg/mL BSA), were mixed with 10 µL of the respective human recombinant enzyme HDAC1 (1.4 ng/µL in assay buffer; BPS Bioscience, Catalog# 50051); HDAC2 (0.95 ng/µl in assay buffer; BPS Bioscience, Catalog# 50052); HDAC6 (3.3 ng/µL in assay buffer; BPS Bioscience, Catalog# 50006) and 25 µL of assay buffer. For HDAC1 and HDAC2 a preincubation step with the inhibitor was performed for 60 min at room temperature, whereas no preincubation was required for HDAC6. Then, 10 µL of the fluorogenic substrate ZMAL (Z-Lys(Ac)-AMC) (75 µM in assay buffer)^68^ was added. After an incubation time of 90 min at 37 °C, 50 µL of 0.4 mg/mL trypsin in trypsin buffer (50 mM Tris-HCl, pH 8.0, 100 mM NaCl) was added, followed by further incubation at 37 °C for 30 min. Fluorescence was measured at an excitation wavelength of 355 nm and an emission wavelength of 460 nm using a FLUOstar OPTIMA microplate reader (BMG Labtech). All compounds were evaluated in duplicates in at least two independent experiments, the 50% inhibitory concentration (IC_50_) was determined by plotting normalized dose-response curves using nonlinear regression (GraphPad Prism 9.0).

#### Cellular HDAC inhibition assay

The cellular HDAC inhibition assay was performed as previously described,^69^ with minor modifications. HeLa^HDAC6-HiBiT^ cells stably expressing the LgBiT protein were seeded at a density of 3 × 10^4^ cells/well in 96-well plates (Greiner Bio-One) using phenol red free DMEM (Life Technologies). The phenol red free medium was supplemented with 10% fetal bovine serum (Sigma Aldrich), 4 mM L-glutamine (PAN Biotech GmbH), 1 mM sodium pyruvate (ThermoFisher Scientific Inc.), 100 IU/mL penicillin, and 0.1 mg/mL streptomycin (PAN Biotech GmbH). Test compounds were first diluted in DMSO and then further diluted in phenol red free cell culture medium. The final DMSO concentration was 0.25%. To test cell permeability, IGEPAL CA-630 (Alfa Aesar, Haverhill, MA, USA) was added at 0.05% for lysed cell assays or omitted for whole cell assays. Compounds were added to the cells, followed by the substrate Boc-Lys(ε-Ac)-AMC (BLD pharma, Reinbeck, Germany) at a final concentration of 300 µM. Cells were incubated for 2 hours at 37 °C and 5% CO_2_ under humidified air. The final assay volume was 50 µL. After incubation, 50 µL of stop solution was added. This solution contained vorinostat 10 µM, 1% IGEPAL CA-630, and 2 mg/mL trypsin in buffer (50 mM Tris-HCl, 137 mM NaCl, 2.7 mM KCl, and 1 mM MgCl_2_). Plates were incubated for another 30 minutes at 37 °C. Fluorescence was measured at an excitation wavelength of 355 nm and an emission wavelength of 460 nm. Data were analyzed using Graph Pad Prism (Graph Pad Prism 9.0) with the 3-parameter logistic equation.

### Molecular docking

For the docking studies the crystal structure of human HDAC6 (chain A) was used (PDB ID: 5EDU).^70^ The protein was prepared using the software Molecular Operating Environment (MOE, Chemical Computing Group, version 2022.02)^71^ by adding hydrogen atoms and building missing loops. Additionally, water molecules, ions (except the catalytic zinc ion) and the co-crystallized ligand were removed from the crystal structure. The ligand−triazole conjugate **1** was drawn with ChemDraw (version 21.0.0.28) and prepared in MOE by generating the 3D structure and energy minimization using AMBER:EHT force field. The docking experiment was performed with GOLD^72^ in combination with the HERMES visualizer (Cambridge Crystallographic Data Centre, version 2022.3.0). The active site was defined by the zinc ion within a radius of 15 Å. All poses were optimized within 50 GA runs and scored with ChemPLP. The early termination option of the docking runs was not allowed. The search efficacy was set to 200%. The docking was carried out with a scaffold match constraint to place the ligand onto a given scaffold location within the active site. The hydroxamate substructure (CONO) of vorinostat in *Danio rerio* HDAC6 (PDB ID: 5EEI)^70^ was used as a template. The docking solutions were visually inspected according to coordination of the zinc ion and a reasonable orientation of the cap groups. The pose depicted in Figure 2A represents the top ranked docking solution out of 50.

### Chemistry

#### General remarks

Chemicals were purchased from BLDpharm, Carl Roth, Fisher Scientific, Sigma Aldrich, Tokyo Chemical Industry and VWR Chemicals. Technical grade solvents were distilled prior to use. For all HPLC purposes, acetonitrile in HPLC-grade quality (HiPerSolv CHROMANORM, VWR) was used. Water was purified with a PURELAB flex® (ELGA VEOLIA). Thin layer chromatography was carried out with pre-coated silica gel (60 F_254_) aluminum sheets from Merck. Detection was performed with UV light at 254 and 360 nm. Solid phase synthesis was processed in PP-reactors equipped with a PE frit (5 mL, 25 µm pore size, MultiSyn Tech GmbH). The synthesis was carried out at r.t. on an orbital shaker (RS-OS 5, Phoenix Instruments GmbH).

*Mass spectrometry*: ESI-MS (LCMS) analyses were carried out on an API 2000 mass spectrometer coupled with an Agilent HPLC HP 1100 using an EC50/2 Nucleodur C18 Gravity 3 μm column or on an Agilent Infinity Lab LC/MSD-system coupled with an Agilent HPLC 1260 Infinity II using an EC50/2 Nucleodur C18 Gravity 3 μm column. The purity of synthesized compounds was determined by HPLC-DAD. HR-ESI-MS spectra were recorded on a Bruker micrOTOF-Qmass spectrometer coupled with a HPLC Dionex UltiMate 3000 or a LTQOrbitrap XL (*Thermo Fisher Scientific Inc.*).

*Nuclear magnetic resonance spectroscopy (NMR)*: NMR spectra were recorded on a Bruker Avance III 600 (600 MHz ^1^H NMR, 151 MHz ^13^C NMR, 565 MHz ^19^F NMR). Chemical shifts are given in parts per million (ppm) referring to the signal centre using the solvent peaks for reference DMSO-*d*_6_ (2.49/39.7). The multiplicity of each signal is reported as singlet (s), doublet (d), triplet (t), multiplet (m) or combinations thereof. Multiplicities and coupling constants are reported as measured and might disagree with the expected values.

*High performance liquid chromatography (HPLC)*: A Thermo Fisher Scientific UltiMate^TM^ 3000 UHPLC system with a Nucleodur 100-5 C18 (250 × 4.6 mm, Macherey Nagel) with a flow rate of 1 mL/min and a temperature of 25 °C with an appropriate gradient was used. For preparative purposes, an AZURA Prep. 500/1000 gradient system with a Nucleodur 110-5 C18 HTec (150 × 32 mm, Macherey Nagel) column and a flow rate of 20 mL/min was used. Detection was implemented with UV absorption measurement at wavelengths of λ = 220 nm and λ = 250 nm. If not stated otherwise, the indicated purity was determined at a wavelength of 250 nm. Bidest. H_2_O with an addition of 0.1% TFA (A) and MeCN (B) were used. The following gradient was used for purification via preparative HPLC: **Method A**: 95% (A) for 5 min equilibration, in 35 min to 95% (B) and 8 min isocratic; **Method B**: 95% (A) for 5 min equilibration, in 5 min to 30% (B), 20 min to 95% (B) and 5 min isocratic.

*Preparation of the preloaded resin* **1**. The synthesis was performed according to our previously published protocols.^23,73^

#### Compounds synthesis and characterization

*2-[6-(Dimethylamino)-3-(dimethyliminio)-3H-xanthen-9-yl]-5-[(3-{4-[(4-{[7-(hydroxyamino)-7-oxoheptyl]carbamoyl}phenoxy)methyl]-1H-1,2,3-triazol-1-yl}propyl)carbamoyl]benzoate. TFA (**4**)*.

HAIR D (**2**)^56^ was coupled with 4-(prop-2-yn-1-yloxy)benzoic acid, which was synthesized according to literature, with our previously published method^73^ to form **3** as an intermediate. After swelling of **3** (180 mg, 0.13 mmol, 1.00 eq.) in DMF (3 mL) for 60 min, the Fmoc-deprotection was performed as described in Feller *et al.*^73^ In parallel, the following solutions were prepared: 5-carboxytetramethylrhodamine-PEG3-azide (81 mg, 0.13 mmol, 1.00 eq.) in DMF (0.10 M), tris[(1-benzyl-4-triazolyl)methyl]amine (TBTA) (17 mg, 0.03 mmol, 0.25 eq.) in DMF (0.10 M), 0.10 M aqueous CuSO_4_*5 H_2_O solution (323 µL, 0.03 mmol, 0.25 eq.) in DMF (0.05 M), 0.20 M aqueous ascorbic acid solution (232 µL, 0.07 mmol, 0.50 eq.) in DMF (0.10 M) and *t*-BuOH (640 µL). After successful Fmoc-deprotection, all prepared solutions were added to the resin, in the order described above, and shaken for 20 h at room temperature. Subsequently, the resin was washed with DMF (5 × 3 mL) and CH_2_Cl_2_ (10 × 3 mL) and dried *in vacuo*. Cleavage according to previously published protocol^73^ using CH_2_Cl_2_/TFA/TIPS, 95/5/5 (*v*/*v*/*v*) and purification by preparative HPLC (Method A) afforded **4** as an amorphous violet powder (67 mg, 71 µmol).

Yield 55%; **^1^H NMR** (600 MHz, DMSO-*d*_6_) δ = 10.32 (s, 1H), 8.94 (t, *J* = 5.6 Hz, 1H), 8.70 (s, 1H), 8.30 (d, *J* = 8.5 Hz, 1H), 8.26 (t, *J* = 5.6 Hz, 1H), 8.20 (s, 1H), 7.80 (d, *J* = 8.5 Hz, 2H), 7.57 (d, *J* = 7.9 Hz, 1H), 7.09 – 7.02 (m, 6H), 6.95 (d, *J* = 2.1 Hz, 2H), 5.18 (s, 2H), 4.53 (t, *J* = 5.2 Hz, 2H), 3.82 (t, *J* = 5.2 Hz, 2H), 3.58 (t, *J* = 5.9 Hz, 2H), 3.57 – 3.53 (m, 2H), 3.54 – 3.47 (m, 8H), 3.31 – 3.18 (m, 14H), 1.93 (t, *J* = 7.4 Hz, 2H), 1.53 – 1.44 (m, 4H), 1.32 – 1.22 (m, 4H). C-NH-O*H* signal could not be detected due to solvent exchange; **^13^C NMR** ^a)^ (151 MHz, DMSO-*d*_6_) δ = 169.1, 165.9, 165.5, 164.7, 160.1, 156.8, 156.7, 142.2, 135.9, 131.3, 131.1, 130.6, 130.4, 129.5, 128.9, 127.2, 125.0, 114.6, 114.1, 112.7, 96.2, 69.7, 69.6, 69.6, 69.5, 68.8, 68.6, 61.2, 49.4, 40.5, 40.1, 32.2, 29.1, 28.3, 26.2, 25.1. The ^13^C NMR signals for carbamoyl benzoate C-1, and 3H-xanthene C-3,4a,6,8a,9,9a,10a could not be detected. (8 carbons); **HRMS (ESI)** *m/z* [M+H]^+^ calcd. for C_50_H_60_N_8_O_11_ 949.4454, found 949.4445; **HPLC** Gradient: after 5 min of equilibration, a linear gradient from 95% A to 5% A in 5 min followed by an isocratic regime of 5% A for 12 min was used), *t*_R_ = 10.80 min, 95.1% purity.

*2-[6-(Dimethylamino)-3-(dimethyliminio)-3H-xanthen-9-yl]-5-[(3-{4-[(4-{[7-(hydroxyamino)-7-oxoheptyl]carbamoyl}phenoxy)methyl]-1H-1,2,3-triazol-1-yl}propyl)carbamoyl]benzoate . TFA (**5**)*.

HAIR D (**2**)^56^ was coupled with 4-(prop-2-yn-1-yloxy)benzoic acid, which was synthesized according to literature, with our previously published method^73^ to form **3** as an intermediate. After swelling of **3** (100 mg, 0.07 mmol, 1.00 eq) in DMF (3 mL) for 60 min, the Fmoc-deprotection was performed as described in Feller *et al*.^73^ In parallel, the following solutions were prepared: 5-carboxytetramethylrhodamine azide (29.4 mg, 0.06 mmol, 0.80 eq.) in DMF (0.10 M), tris[(1-benzyl-4-triazolyl)methyl]amine (TBTA) (9.5 mg, 0.02 mmol, 0.25 eq.) in DMF (0.10 M), 0.10 M aqueous CuSO_4_*5 H_2_O solution (179 µL, 0.02 mmol, 0.25 eq.) in DMF (0.05 M), 0.20 M aqueous ascorbic acid solution (179 µL, 0.04 mmol, 0.50 eq.) in DMF (0.10 M) and *t*-BuOH (355 µL). After successful Fmoc-deprotection, all prepared solutions were added to the resin, in the order described above, and shaken for 20 h at room temperature. Subsequently, the resin was washed with DMF (5 × 3 mL) and CH_2_Cl_2_ (10 × 3 mL) and dried *in vacuo*. Cleavage according to previously published protocol^73^, using CH_2_Cl_2_/TFA/TIPS, 95/5/5 (*v*/*v*/*v*) and purification by preparative HPLC (Method A) afforded **5** as an amorphous violet powder (22.3 mg, 27 µmol).

Yield 47%; **^1^H NMR** (600 MHz, DMSO-*d*_6_) δ = 10.31 (s, 1H), 8.96 (t, *J* = 5.6 Hz, 1H), 8.69 (s, 1H), 8.35 – 8.24 (m, 3H), 7.84 – 7.78 (m, 2H), 7.59 (d, *J* = 7.9 Hz, 1H), 7.15 – 6.89 (m, 8H), 5.21 (s, 2H), 4.50 (t, *J* = 7.0 Hz, 2H), 3.40 – 3.37 (m, 2H), 3.32 – 3.16 (m, 14H), 2.21 – 2.12 (m, 2H), 1.93 (t, *J* = 7.4 Hz, 2H), 1.53 – 1.44 (m, 4H), 1.32 – 1.22 (m, 4H), C-NH-O*H* signal could not be detected due to solvent exchange; **^13^C NMR** (151 MHz, DMSO-*d*_6_) δ = 169.1, 165.9, 165.5, 164.8, 160.1, 156.7, 142.3, 135.9, 131.2, 130.5, 129.5, 128.9, 127.3, 124.7, 114.6, 114.1, 112.7, 96.3, 61.3, 47.5, 40.5, 40.1, 36.8, 32.2, 29.7, 29.1, 28.3, 26.2, 25.1. The ^13^C NMR signals for carbamoyl benzoate C-1,5, -COOH, and 3*H*-xanthene C-3,4a,6,8a,9,9a,10a could not be detected. (10 carbons). **HRMS (ESI)** *m/z* [M+H]^+^ calcd. for C_45_H_50_N_8_O_8_ 831.3824, found 831.3821; **HPLC** (gradient: After 5 min of equilibration, a linear gradient from 95% A to 5% A in 5 min followed by an isocratic regime of 5% A for 12 min was used), *t*_R_ = 10.83 min, 99.0% purity.

*Methyl 4-((2-acetyl-1-phenylhydrazineyl)methyl)-3-fluorobenzoate (6)*

2-Acetyl-1-phenylhydrazine (1.00 g, 6.66 mmol, 1.00 eq.) and DIPEA (1.31 mL, 7.52 mmol, 1.13 eq.) were dissolved in MeCN (8 mL). Methyl 4-bromomethylbenzoate (1.86 g, 7.52 mmol, 1.13 eq.) was added and the reaction was allowed to stir at 95 °C for 18 h. The reaction mixture was evaporated *in vacuo* and the residue was dissolved in CH_2_Cl_2_, washed with distilled H_2_O (2 × 40 mL) and once with brine (40 mL), dried over Na_2_SO_4_, filtered, and evaporated *in vacuo*. The crude product was purified on silica (EtOAc in *c*-Hex 40 to 60%) providing **6** as a beige solid (1.9 g, yield: 92%). *R_f_* (c-Hex/EtOAc 4:6 *v/v)*: 0.49. **^1^H NMR** (500 MHz, CDCl_3_; chemical shifts reported for both rotamers) δ 7.81 – 7.75 (m, 3H), 7.75 – 7.69 (m, 2H), 7.35 – 7.29 (m, 3H), 7.25 – 7.20 (m, 2H), 7.00 – 6.95 (m, 3H), 6.89 – 6.80 (m, 3H), 4.81 (s, 3H), 3.92 (s, 3H), 3.91 (s, 3H), 2.00 (s, 3H), 1.83 (s, 3H). **^13^C NMR** (126 MHz, CDCl_3_) δ 176.0, 169.3, 165.9, 165.7, 161.9, 161.6, 159.9, 159.6, 148.7, 148.1, 132.3, 131.6, 131.0, 130.4, 129.8, 129.5, 128.1, 128.0, 125.6, 121.8, 120.3, 114.4, 113.0, 53.2, 51.2, 21.0, 19.2. The ^13^C NMR peak number exceeds the expected one due to the presence of rotamers. **^19^F NMR** (471 MHz, CDCl_3_) δ -116.75 (dd, *J* = 10.3, 7.3 Hz), -117.53 (dd, *J* = 10.4, 7.2 Hz). **LC-MS (ESI)** *m/z* [M+H]^+^ calcd. for C_17_H_17_FN_2_O_3_ 317,1 found 317.2.

*3-Fluoro-4-((1-phenylhydrazineyl)methyl)benzoic acid . HCl (**7**)*

AcOH (4.2 mL) and HCl 6 M (12.5 mL) was added to compound **6** (1.93 g, 6.11 mmol, 1.00 eq.) and the reaction was allowed to stir for 2 h at 110 °C. The precipitate was filtered and dried *in vacuo*. Compound **7** was obtained as white powder and used with no further purification (1.35 g, yield: 74%). **^1^H NMR** (600 MHz, DMSO-*d*_6_) δ 10.51 (s, 1H), 7.73 (dd, *J* = 7.9, 1.6 Hz, 1H), 7.64 (dd, *J* = 10.4, 1.6 Hz, 1H), 7.52 (t, *J* = 7.6 Hz, 1H), 7.38 – 7.32 (m, 2H), 7.21 – 7.18 (m, 2H), 7.13 (t, *J* = 7.4 Hz, 1H), 4.77 (s, 2H); **^13^C NMR** (151 MHz, DMSO-*d*_6_) δ 166.0, 160.9, 159.3, 146.5, 132.7, 131.3, 129.1, 127.2, 125.2, 124.4, 119.1, 116.0, 115.8, 53.0. **^19^F NMR** (565 MHz, DMSO-*d*_6_) δ -115.74 – -116.45 (m).

*4-((2-(((9H-Fluoren-9-yl)methoxy)carbonyl)-1,2,3,4-tetrahydro-5H-pyrido[4,3-b]indol-5-yl)methyl)-3-fluorobenzoic acid (**8**)*

Compound **7** (1.35 g, 4.42 mmol, 1.00 eq.) was dissolved in AcOH (8 mL) and 1-Fmoc-4-piperidone (1.45 g, 4.51 mmol, 1.02 eq.) was added. The reaction was allowed to stir for 2 h at 110°C. After completion of the reaction, the mixture was evaporated *in vacuo*, dissolved in EtOAc (40 ml) and washed with distilled H_2_O (3 × 40 mL) and once with brine (20 mL). The organic layer was dried over with Na_2_SO_4_, filtered, and dried *in vacuo*. The crude product was purified by silica gel chromatography (MeOH in CH_2_CL_2_ 0 to 10% + 0.1% formic acid *v*/*v*), providing compound **8** as a beige solid (1.83 g, yield: 76%). *R_f_* (CH_2_Cl_2_/MeOH/formic acid 94.95:4.95:0.1 *v/v/v*): 0.55. **^1^H NMR** (600 MHz, DMSO-*d*_6_) δ 7.92 – 7.76 (m, 2H), 7.64 (d, *J* = 40.5 Hz, 4H), 7.44 (dd, *J* = 29.9, 8.0 Hz, 2H), 7.36 (s, 2H), 7.30 – 7.16 (m, 2H), 7.07 (d, *J* = 34.4 Hz, 2H), 6.76 (t, *J* = 7.7 Hz, 1H), 5.47 (s, 2H), 4.55 (d, *J* = 16.4 Hz, 2H), 4.42 (s, 2H), 4.30 (t, *J* = 6.4 Hz, 1H), 3.72 (s, 1H), 3.62 (d, *J* = 17.5 Hz, 1H), 2.71 (s, 1H), 2.56 (s, 1H); **^13^C NMR** (151 MHz, DMSO-*d*_6_) δ 170.3, 163.1 (d, *J* = 886.9 Hz), 158.5, 143.9, 140.8, 136.3, 133.7 (d, J = 39.5 Hz), 133.5, 132.7, 129.7 (d, J = 14.9 Hz), 128.5, 127.5, 127.0, 125.6, 124.9, 124.8, 121.2, 120.0, 119.3, 117.5, 115.9, 115.8, 109.6, 106.6, 66.6, 59.7, 46.7, 41.0, 20.7. The ^13^C NMR peak number exceeds the expected number due to the presence of rotamers. **^19^F NMR** (565 MHz, DMSO-*d*_6_) δ -117.05 (d, *J* = 69.4 Hz). **LC-MS (ESI)** *m/z* [M+H]^+^ calcd. for C_34_H_27_FN_2_O_4_ 547,195 found 547.300.

*4-((2-(8-(2-(4-(2,6-Dioxopiperidin-3-yl)phenoxy)acetamido)octanoyl)-1,2,3,4-tetrahydro-5H-pyrido[4,3-b]indol-5-yl)methyl)-3-fluoro-N-hydroxybenzamide* (MM-47)

The resin **12**^23,56^ (330 mg, 0.528 mmol) was swelled in DMF (1 mL) for 5 min and coupled with a solution of **8** (595 mg, 1.056 mmol, 2.00 eq.), HATU (406 mg, 1.056 mmol, 2 eq.), HOBt . H_2_O (202 mg, 1.056 mmol, 2.00 eq.), and DIPEA (279 µL, 1.584 mmol, 3.00 eq.) in DMF (1.3 mL) for 48 h at room temperature. Subsequently, the resin was washed with DMF (5 × 5 mL) and CH_2_Cl_2_ (10 × 5 mL). The completion of the reaction was confirmed *via* TNBS test, conducted according to manufacturer’s protocol, and by HPLC analysis after test cleavage (cleavage solution: TFA 5% in CH_2_Cl_2_ *v*/*v*). The Fmoc-deprotection was performed as described in Feller *et al.*^73^ and the resin was washed with DMF (5 × 5 mL), MeOH (5 × 5 mL), and DMF (5 × 5 mL). In parallel, 8-(Fmoc-amino)octanoic acid (136 mg, 0.345 mmol, 2.00 eq.), HATU (132 mg, 0.345 mmol, 2.00 eq.), HOBt . H_2_O (66 mg, 0.345 mmol, 3.00 eq.), and DIPEA (91 µL, 0.517 mmol, 3 eq.) were dissolved in DMF (500 µL) and stirred for 5 min. The solution was added to the resin and the amide coupling was performed for 18 h at room temperature. After completion of the reaction, the resin was washed with DMF (5 × 5 mL) and CH_2_Cl_2_ (10 × 5 mL). The completion of the reaction was confirmed *via* TNBS test, conducted according to manufacturer’s protocol, and by HPLC analysis after test cleavage (cleavage solution: TFA 5% in CH_2_Cl_2_ *v*/*v*). Subsequently, Fmoc-deprotection was performed on resin **14** and afterwards the deprotected resin was washed with DMF (5 × 5 mL), MeOH (5 × 5 mL) and DMF (5 × 5 mL). The final coupling was performed using the deprotected resin **14** (82.5 mg, 0.042 mmol, 1.0 eq.) and a solution of 2-(4-(2,4-dioxocyclohexyl)phenoxy)acetic acid (**11**, 10 mg, 0.035 mmol, 0.85 eq.), HATU (13 mg, 0,035 mmol, 0.85 eq.) HOBt . H_2_O (5 mg, 0.035 mmol, 0.85 eq.), and DIPEA (12 µL, 0.070 mmol, 1.65 eq.) in DMF (100 µL) for 48 h at room temperature. After 48 h, a second cycle of amide coupling was performed, using the same conditions. Subsequently, the resin was washed with DMF (5 × 3 mL), CH_2_Cl_2_ (10 × 3 mL), and dried *in vacuo*. Cleavage according to previously published protocol^56^, using CH_2_Cl_2_/TFA, 95/5 (*v*/*v*) and purification by preparative HPLC (Method B) afforded **MM-47** as an amorphous light yellow solid. (8 mg, 11 µmol). Yield: 26%; **^1^H NMR** (600 MHz, DMSO-*d*_6_) δ 11.23 (s, 1H), 10.77 (s, 1H), 8.01 (q, *J* = 6.0 Hz, 1H), 7.55 (ddd, *J* = 11.0, 5.1, 1.6 Hz, 1H), 7.50 (dd, *J* = 18.6, 7.8 Hz, 1H), 7.45 (ddd, *J* = 7.4, 5.5, 1.6 Hz, 1H), 7.41 – 7.37 (m, 1H), 7.16 – 7.11 (m, 2H), 7.09 (t, *J* = 7.6 Hz, 1H), 7.03 (td, *J* = 7.4, 3.4 Hz, 1H), 6.90 (d, *J* = 8.3 Hz, 2H), 6.80 – 6.73 (m, 1H), 5.45 (d, *J* = 2.6 Hz, 2H), 4.68 (d, *J* = 17.1 Hz, 2H), 4.43 (s, 2H), 3.85^#^ (t, *J* = 5.8 Hz, 2H), 3.79 (dq, *J* = 11.5, 5.2 Hz, 5H, partially overlaid by solvent signal), 3.10 (p, *J* = 7.2 Hz, 2H), 2.81 (d, *J* = 11.7 Hz, 1H), 2.72 (t, *J* = 5.7 Hz, 1H), 2.64 (ddd, *J* = 17.1, 11.7, 5.2 Hz, 1H), 2.42 (q, *J* = 7.4 Hz, 2H), 2.15 (qd, *J* = 11.9, 4.3 Hz, 1H), 2.00 (dq, *J* = 13.7, 4.9 Hz, 1H), 1.51 (h, *J* = 7.7 Hz, 2H), 1.42 (dq, *J* = 15.2, 7.7 Hz, 2H), 1.31 – 1.16 (m, 6H); **^13^C NMR**^a)^ (151 MHz, DMSO-*d*_6_) δ 174.3, 173.4, 171.4, 171.2, 167.4, 162.6, 159.3 (d, J = 246.2 Hz), 156.7, 136.3, 136.2, 134.3, 134.0, 133.7, 131.7, 129.5, 128.5, 128.1, 125.1, 124.8, 123.1, 121.1, 119.2, 119.1, 117.7, 117.6, 114.5, 113.9, 113.8, 109.6, 106.9, 106.7, 67.1, 46.5, 42.4, 42.0, 40.1, 38.8, 38.6, 38.2, 38.1, 32.9, 32.5, 31.3, 29.0, 28.7, 28.7, 28.5, 26.2, 26.0, 25.4, 24.8, 24.7, 22.7, 21.7. The ^13^C NMR peak number exceeds the expected number due to the presence of rotamers. **^19^F NMR** (565 MHz, DMSO), -116.99 (dt, *J* = 88.0, 9.7 Hz). **HRMS (ESI)** *m/z* [M+H]^+^ calcd. For C_40_H_44_FN_5_O_7_ 725.3225, found 726.3298; **HPLC** (gradient: after 5 min of equilibration, a linear gradient from 95% (A) to 5% (A) in 10 min followed by an isocratic regime of 5% (A) for 12 min was used), *t*_R_ = 14.68 min, 96.1% purity.

## ASSOCIATED CONTENT

### Supporting Information

Supplementary Figures and Schemes; ^1^H, ^13^C, and ^19^F NMR spectra; HPLC data of newly synthesized compounds (PDF)

## Author Contributions

The manuscript was written through contributions of all authors. All authors have given approval to the final version.

## ACKNOWLEDGMENTS

Experimental support by Svenja Henze is gratefully acknowledged. The authors thank Dr. Tim Keuler, Dr. Beate König, Dr. Aleša Bricelj, and Dr. Christian Steinebach for providing certain PROTACs and PROTAC building blocks. Figure 1, Figure 3, Figure 5, and the TOC graphic were created using Biorender.com

## ABBREVIATIONS

ATAT1: *N*-acetyltransferase
AUC: area under the curve
BRET: bioluminescence resonance energy transfer
CD: catalytic domain
CDCl_3_: chloroform-*d*
CRBN: cereblon
DCM: dichloromethane
DFMO: difluoromethyl-1,3,4-oxadiazole
DIPEA: *N*,*N*-diisopropylethylamine
Dmax: maximal degradation
Dmax_50_: half maximal degradation at the timepoint of Dmax
DMB: dynein motor binding domain
DMF: *N*,*N*-dimethylformamide
DMSO: dimethylsulfoxide
EMA: European Medicines Agency
EtOAc: ethyl acetate
HAT: histone acetyltransferases
HATU: hexafluorophosphate azabenzotriazole tetramethyl uranium
HDAC: histone deacetylase
HDACi: histone deacetylase inhibitor(s)
HOBt*H_2_O: hydroxybenzotriazole hydrate
HSF-1: heat shock factor protein 1
Hsp90: heat shock protein 90
IC_50_: half maximal inhibitory concentration
MeOH: methanol; min, minutes
NAD^+^: nicotinamide adenine dinucleotide
NES: nuclear export signal
NLS: nuclear localization signal
Nluc: nanoluciferase
PDB: protein data bank
POI: protein of interest
PROTAC: PROteolysis TArgeting Chimeras
RLU: relative light units
rt: room temperature
SD: standard deviation
SEM: standard error of the mean
SE14: serine-glutamate tetradecapeptide repeat
SIRT: sirtuin
*t*-BuOH: *tert*-butanol
TAMRA: tetramethylrhodamine
TBTA: tris(benzyltriazolylmethyl)amine
TFA: trifluoroacetic acid
TFMO: trifluoromethyl-1,3,4-oxadiazole
TIPS: triisopropylsilane
TPD: targeted protein degradation
UPS: ubiquitin-proteasome system
U.S. FDA: United States Food and Drug Administration
VHL: von Hippel-Lindau
ZnF-UBP: ubiquitin-binding zinc finger domain.

## Notes

The authors declare no competing financial interest.

