## Supporting Information for "Target engagement studies and kinetic live-cell degradation assays enable the systematic characterization of HDAC6 PROTACs"

### Table of Contents

1 SUPPLEMENTAL FIGURES AND SCHEMES ........ S3

2 NMR SPECTRA ........ S7

3 HPLC CHROMATOGRAMS ........ S11

4 REFERENCES ........ S13

1 SUPPLEMENTAL FIGURES AND SCHEMES

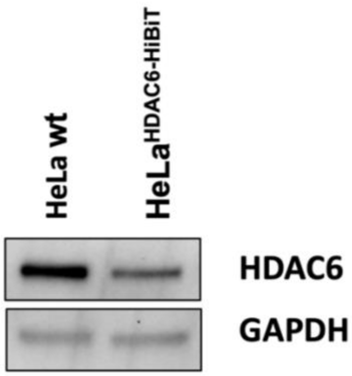


**Figure S1:** Immunoblot analysis of the HDAC6 expression levels of HeLa wild-type and HeLa$^{HDAC6\text{-}HiBiT}$ cells. GAPDH was used as loading control (representative image, n = 2 independent replicates).

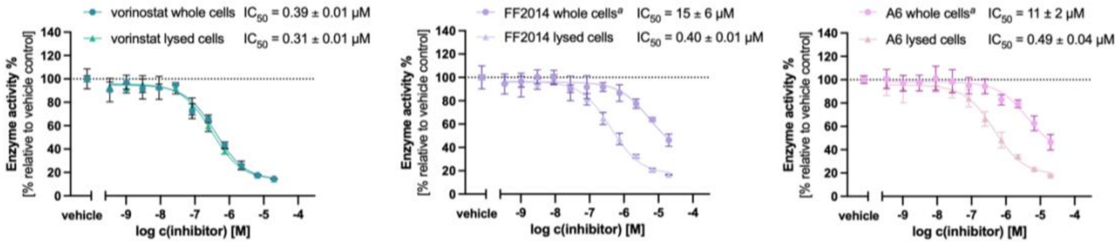


**Figure S2:** Cell permeability study in whole and lysed HeLa$^{HDAC6\text{-}HiBiT}$ cells using a cellular HDAC inhibition assay. Cells were treated with increasing concentrations of the respective compounds. (representative image, mean ± SD, duplicate measurements, n = 2). [a]Curves were adjusted to vorinostat 20 µM.

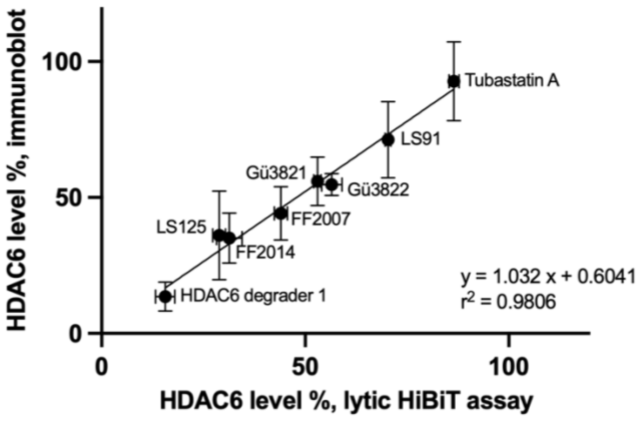


**Figure S3.** Linear regression of HDAC6 level in HeLa$^{HDAC6\text{-}HiBiT}$ cells stably expressing the LgBiT protein determined with HiBiT lytic detection and immunoblot analysis after incubation with compounds at 1 μM for 24 h (mean ± SEM, n = 3).

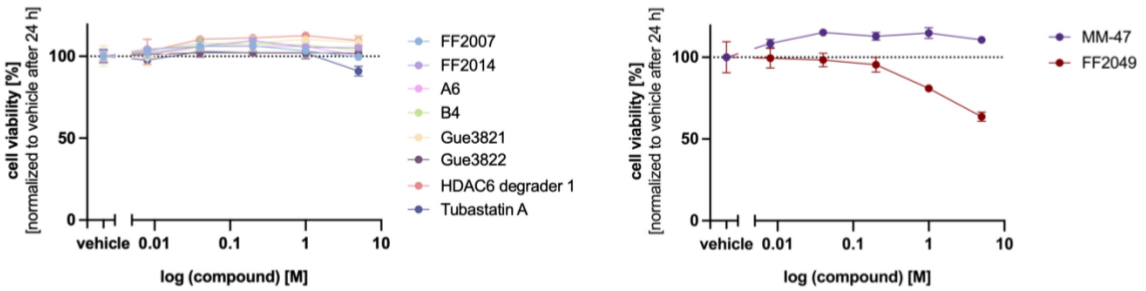


**Figure S4.** Cell viability of HeLa$^{HDAC6\text{-}HiBiT}$ cells stably expressing the LgBiT protein after incubation with PROTACs or tubastatin A in the live cell kinetic HiBiT detection assay for 24 h (representative images, mean ± SD, triplicate measurement, n ≥ 2).

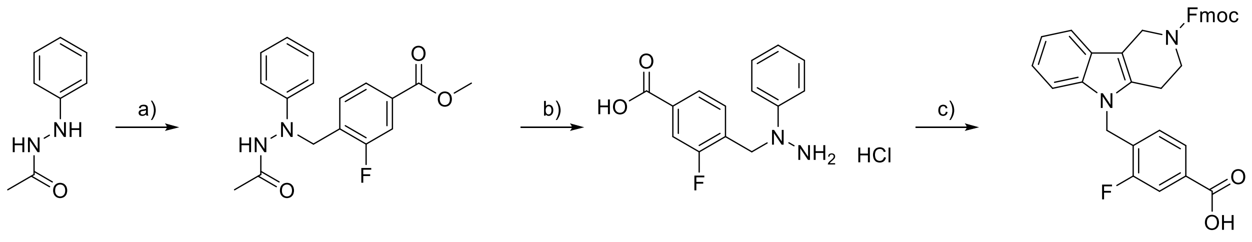


6 7 8

**Scheme S1.** Synthesis of the Fmoc-protected tubastatin A derivative **8** according to literature known protocols[1] with slight modifications. *Reagents and conditions*: a) Methyl 4-(bromomethyl)-3-fluorobenzoate, DIPEA, MeCN, 95 °C, 18 h. b) 4 M HCl, AcOH, 110 °C, 2 h. c) *N*-Fmoc-4-piperidone, AcOH, 110 °C, 2 h.

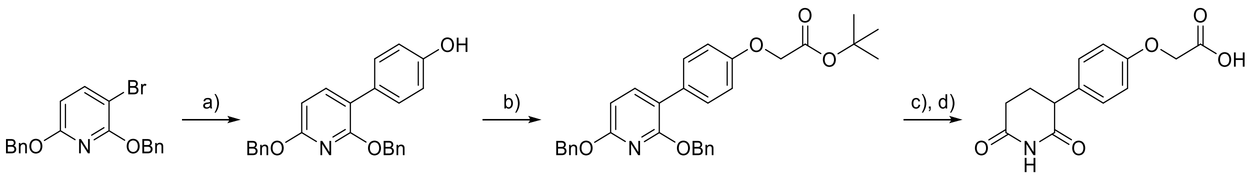


9 10 11

**Scheme S2.** Synthesis of the phenyl-glutarimide based ligand **11** according to a literature known protocol.[2] *Reagents and conditions*: a) 4-hydroxybenzeneboronic acid, $PdCl_2$(dppf)-DCM, $K_3PO_4$, 1,4-dioxane, $H_2O$, 100 °C, 16 h; b) *tert*-butyl bromoacetate, $K_2CO_3$, DMF, 23 °C, 3 h; c) $H_2$, 10% Pd/C, EtOH, 23 °C, 16 h; d) TFA, DCM, 23 °C, 16 h.

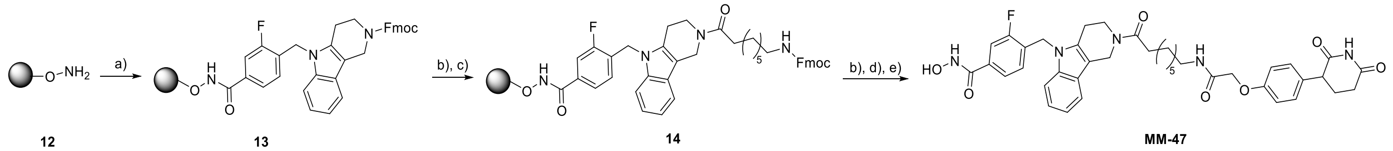


12 13 14 MM-47

**Scheme S3.** Solid-phase synthesis of the HDAC6 degrader **MM-47**. *Reagents and conditions*: a) **8**, HATU, HOBt · $H_2O$ , DIPEA, DMF, 23 °C, 48 h; b) piperidine 20% in DMF, 23 °C, 2 x 5 min;

c) 8-(Fmoc-amino)octanoic acid, HATU, HOBt · $H_2O$, DIPEA, DMF, 23 °C, 18 h; d) **11**, HATU, HOBt · $H_2O$, DIPEA, DMF, 23 °C, 48 h; (e) TFA, DCM, 23 °C, 1 h.

2 NMR SPECTRA

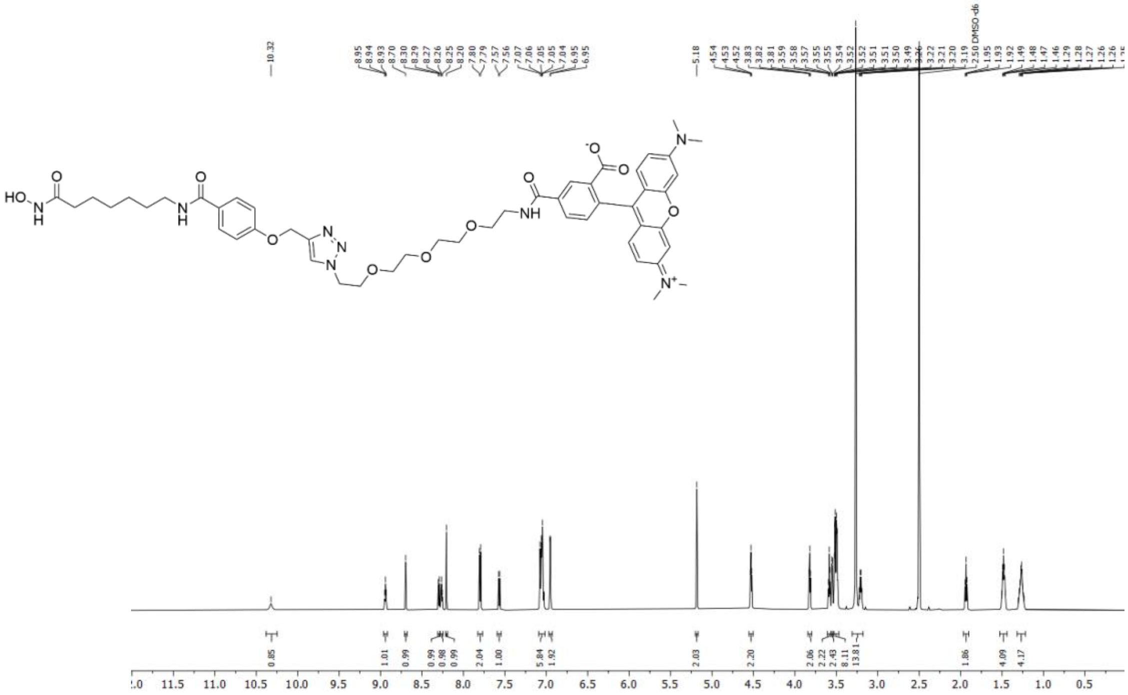

**$^{1}$H NMR** spectrum of **4** (600 MHz, DMSO-$d_6$).

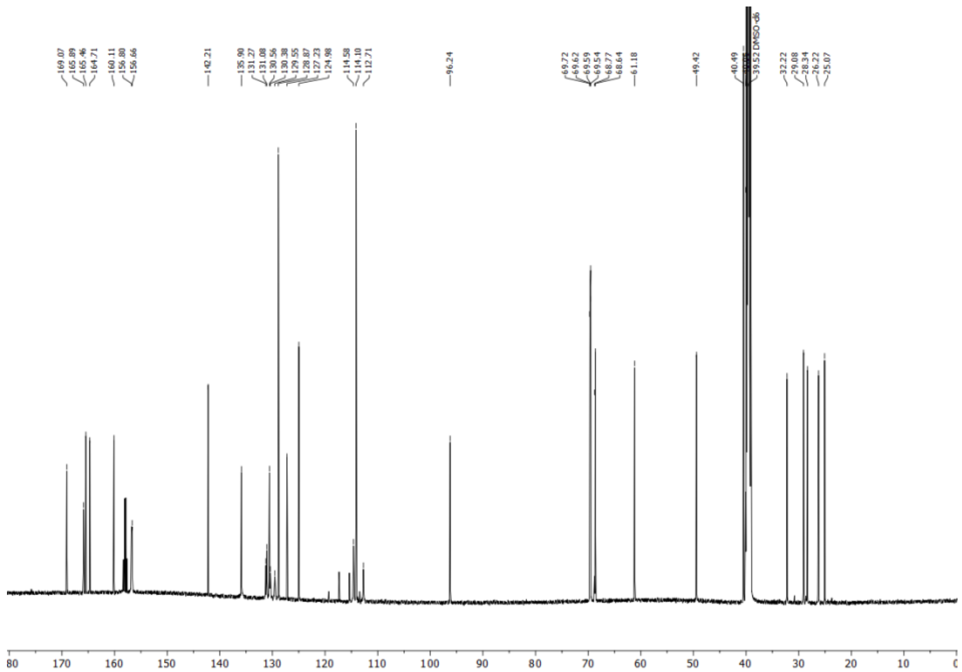

**$^{13}$C NMR** spectrum of **4** (151 MHz, DMSO-$d_6$).

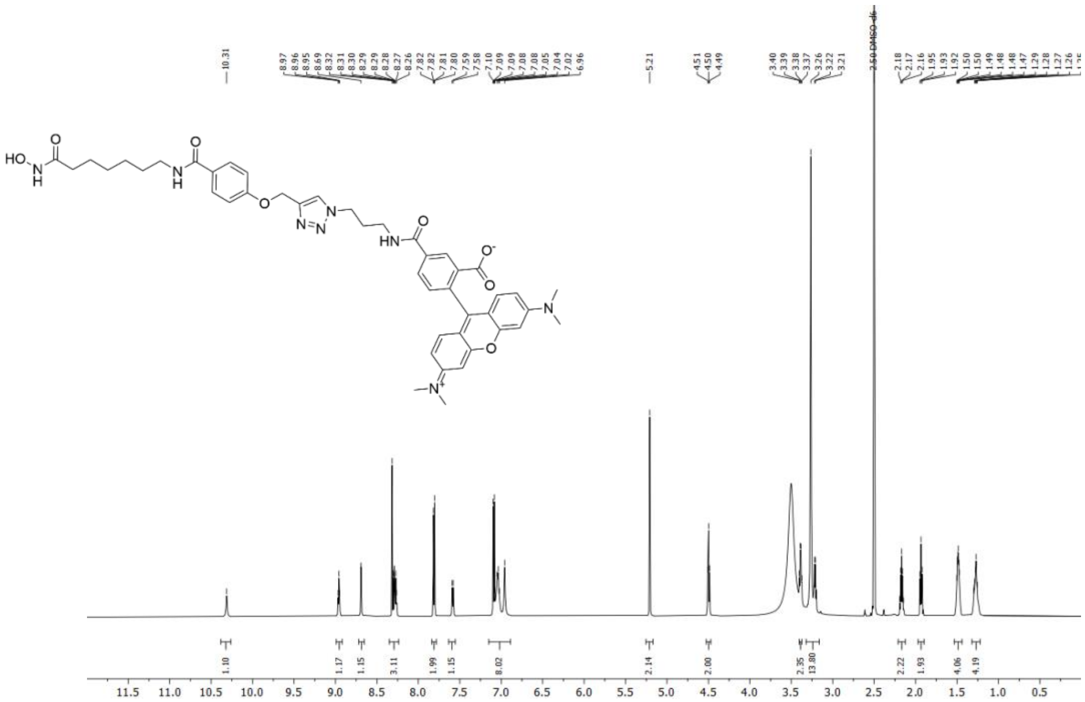


**$^{1}$H NMR** spectrum of **5** (600 MHz, DMSO-$d_6$).

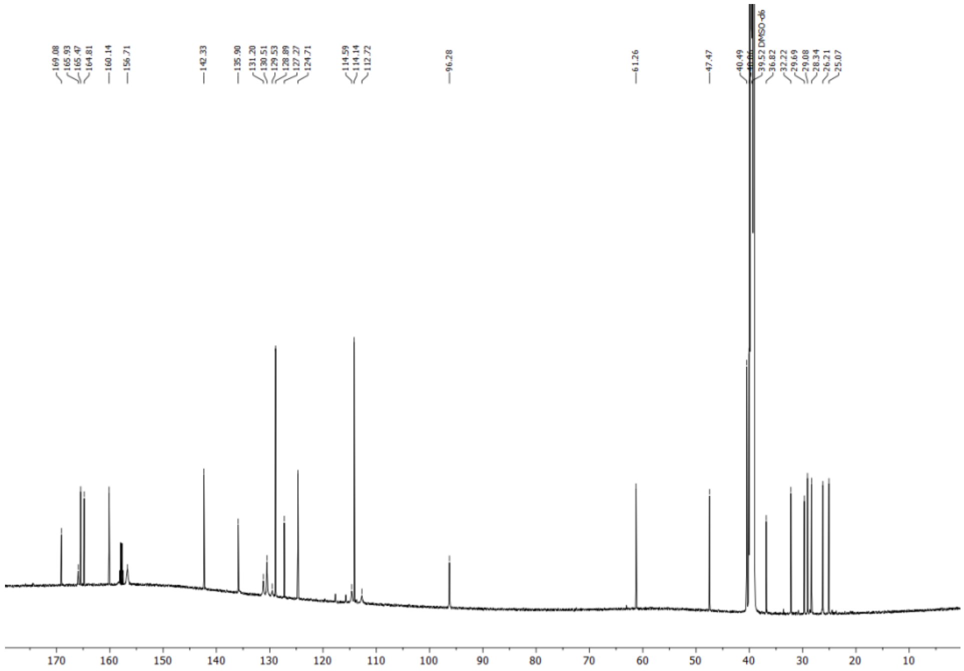


**$^{13}$C NMR** spectrum of **5** (151 MHz, DMSO-$d_6$).

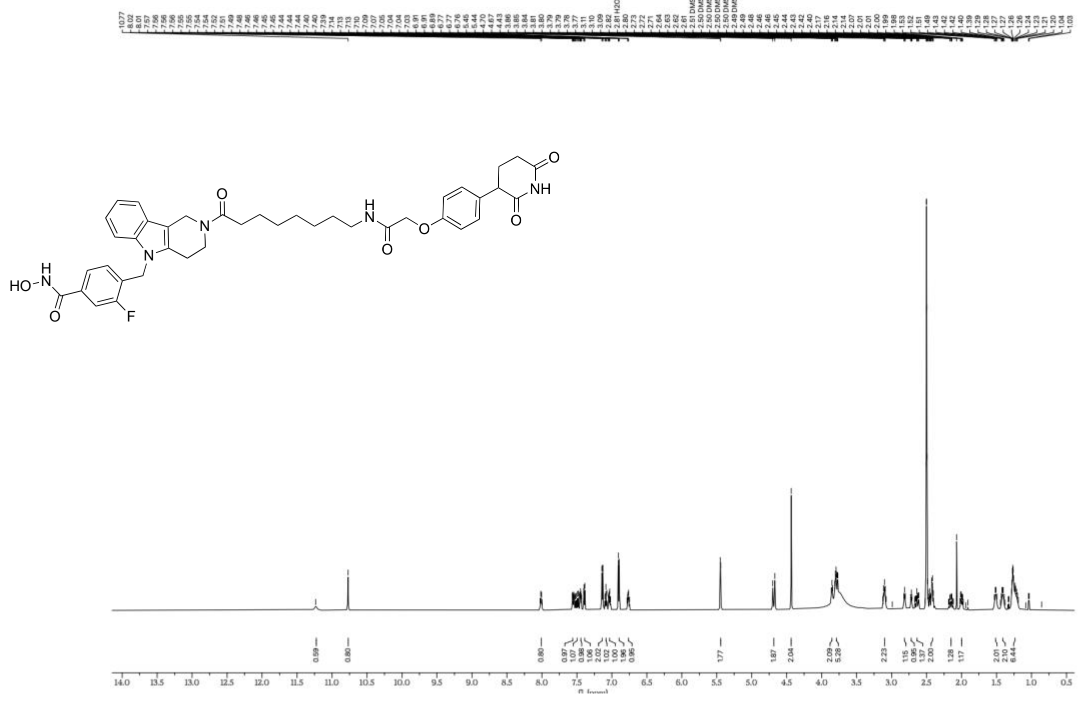

**$^{1}$H NMR** spectrum of **MM-47** (600 MHz, DMSO-$d_6$).

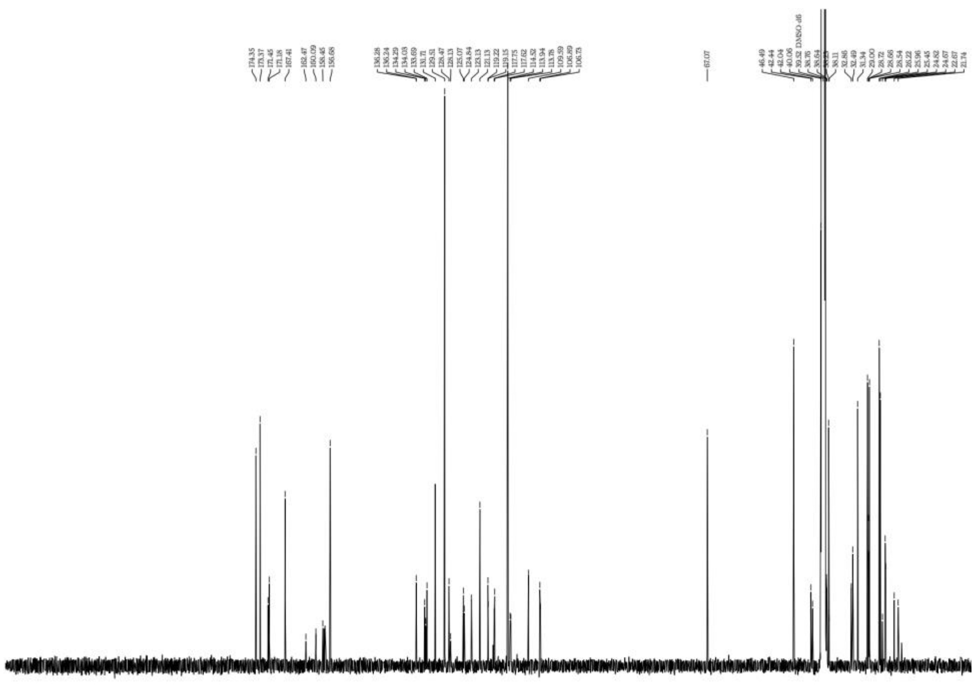

**$^{13}$C NMR** spectrum of **MM-47** (151 MHz, DMSO-$d_6$).

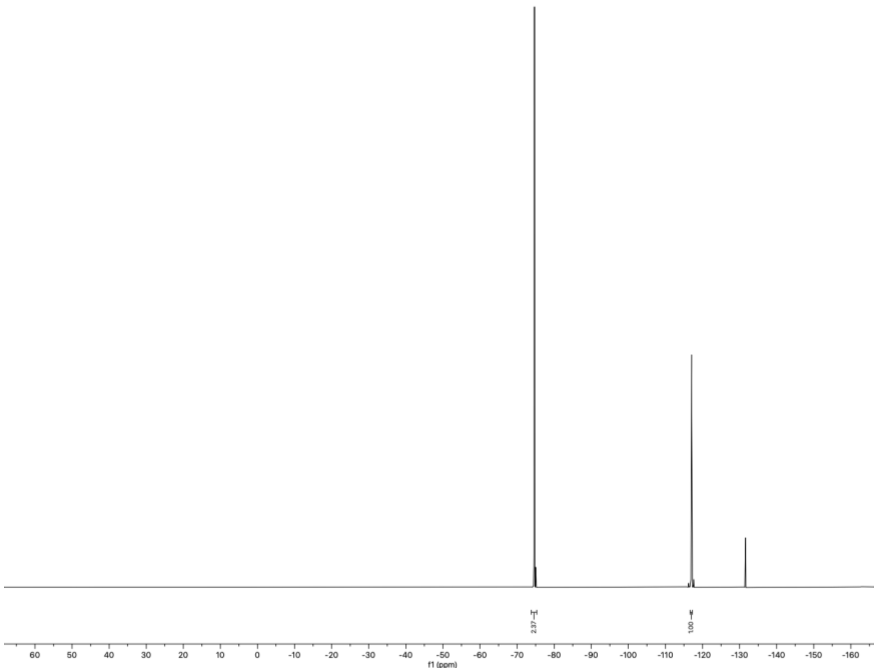


**$^{19}$F NMR** spectrum of **MM-47** (565 MHz, DMSO-$d_6$).

### 3 HPLC CHROMATOGRAMS

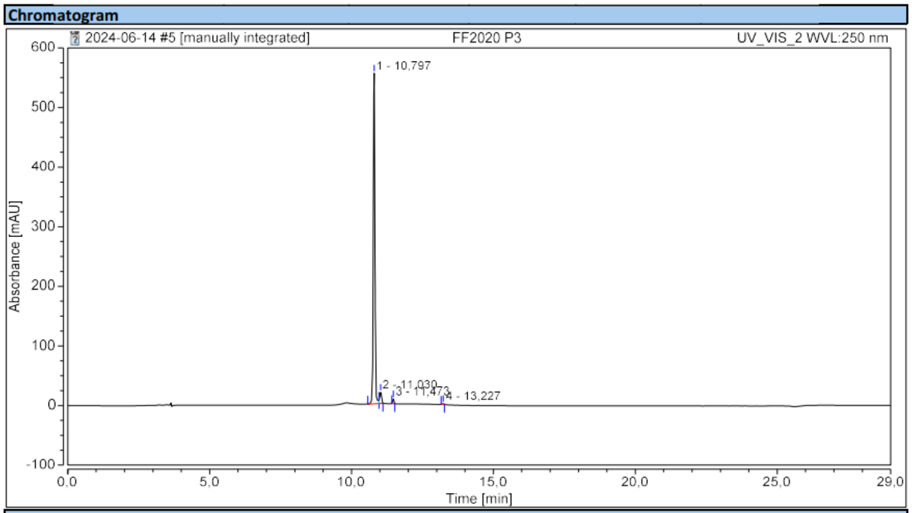


**Integration Results**

| No. | Peak Name | Retention Time min | Area mAU*min | Height mAU | Relative Area % | Relative Height % | Amount n.a. |
|---|---|---|---|---|---|---|---|
| 1 | | 10,797 | 36,160 | 554,493 | 95,13 | 95,29 | n.a. |
| 2 | | 11,030 | 1,357 | 18,512 | 3,57 | 3,18 | n.a. |
| 3 | | 11,473 | 0,418 | 7,785 | 1,10 | 1,34 | n.a. |
| 4 | | 13,227 | 0,077 | 1,099 | 0,20 | 0,19 | n.a. |
| **Total:** | | | **38,012** | **581,889** | **100,00** | **100,00** | |

**HPLC** chromatogram of **4** (purity: 95.1%).

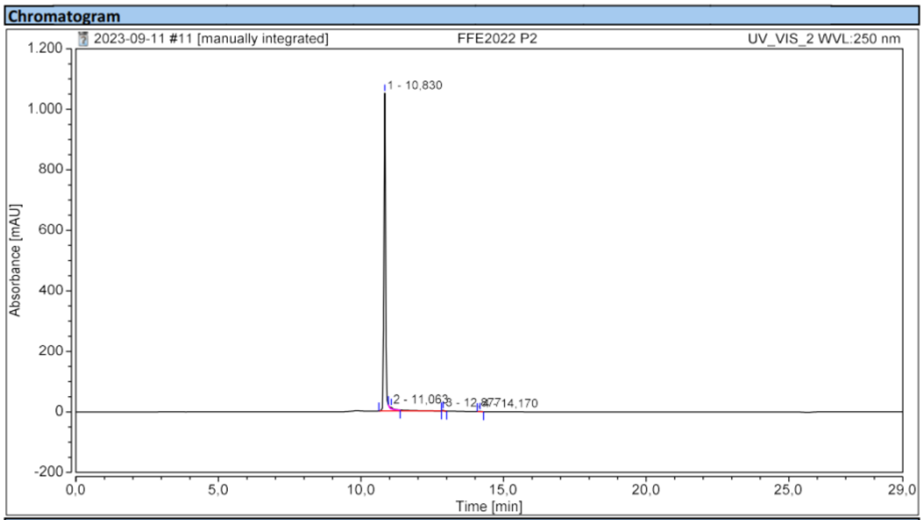


**Integration Results**

| No. | Peak Name | Retention Time min | Area mAU*min | Height mAU | Relative Area % | Relative Height % | Amount n.a. |
|---|---|---|---|---|---|---|---|
| 1 | | 10,830 | 75,723 | 1050,985 | 99,01 | 99,14 | n.a. |
| 2 | | 11,063 | 0,509 | 5,626 | 0,67 | 0,53 | n.a. |
| 3 | | 12,877 | 0,160 | 2,395 | 0,21 | 0,23 | n.a. |
| 4 | | 14,170 | 0,092 | 1,084 | 0,12 | 0,10 | n.a. |
| **Total:** | | | **76,484** | **1060,091** | **100,00** | **100,00** | |

**HPLC** chromatogram of **5** (purity: 99.0%).

Chromatogram

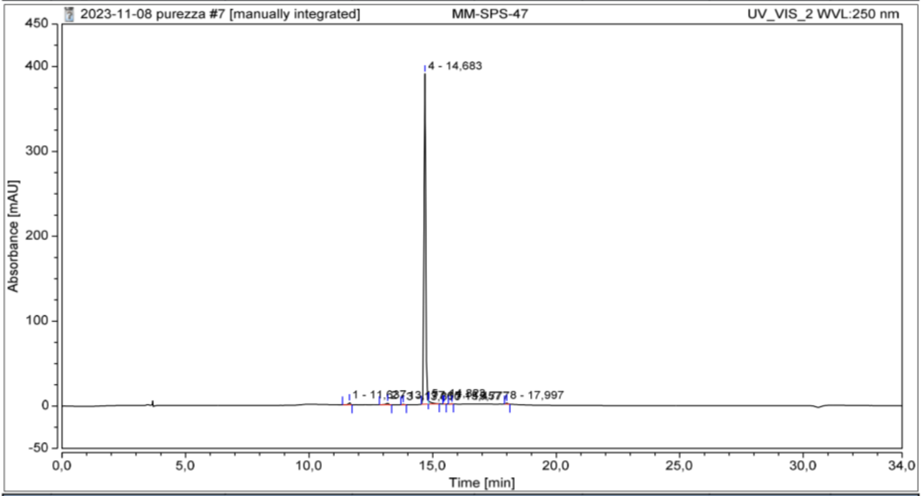


Integration Results

| No. | Peak Name | Retention Time min | Area mAU*min | Height mAU | Relative Area % | Relative Height % | Amount n.a. |
|---|---|---|---|---|---|---|---|
| 1 | | 11,637 | 0,240 | 2,702 | 0,74 | 0,67 | n.a. |
| 2 | | 13,177 | 0,271 | 2,123 | 0,83 | 0,53 | n.a. |
| 3 | | 13,810 | 0,057 | 0,650 | 0,17 | 0,16 | n.a. |
| 4 | | 14,683 | 31,323 | 389,383 | 96,15 | 97,17 | n.a. |
| 5 | | 14,823 | 0,536 | 3,789 | 1,65 | 0,95 | n.a. |
| 6 | | 15,457 | 0,011 | 0,161 | 0,03 | 0,04 | n.a. |
| 7 | | 15,777 | 0,029 | 0,415 | 0,09 | 0,10 | n.a. |
| 8 | | 17,997 | 0,111 | 1,491 | 0,34 | 0,37 | n.a. |
| **Total:** | | | **32,578** | **400,713** | **100,00** | **100,00** | |

**HPLC** chromatogram of **MM-47** (purity: 96.1%).
